# A Novel Therapeutic Approach using CXCR3 Blockade to Treat Immune Checkpoint Inhibitor-mediated Myocarditis

**DOI:** 10.1101/2024.01.30.576279

**Authors:** Yuhsin Vivian Huang, Daniel Lee, Yin Sun, Harrison Chou, Bruce Xu, Zachary Lin, Corynn Branche, Abraham Bayer, Sarah Waliany, Joel Neal, Heather Wakelee, Ronald Witteles, Patricia Nguyen, Edward Graves, Pilar Alcaide, Gerald J. Berry, Sean M. Wu, Han Zhu

## Abstract

**Background:** Immune checkpoint inhibitors (ICIs) are successful in treating many cancers but may cause immune-related adverse events. ICI-mediated myocarditis has a high fatality rate of up to 40%, with severe cardiovascular consequences. Targeted therapies for ICI myocarditis are currently lacking.

**Methods:** We used a genetic mouse model of *PD-1* deletion (*MRL/Pdcd1-/-*) along with a novel drug-treated ICI myocarditis mouse model to recapitulate the disease phenotype. We performed single-cell RNA-sequencing (scRNAseq), single-cell T-cell receptor sequencing (scTCR-seq), and cellular indexing of transcriptomes and epitopes (CITE-seq) on immune cells isolated from *MRL* and *MRL/Pdcd1-/-* mice at serial timepoints. We assessed the impact of macrophage deletion in *MRL/Pdcd1-/-* mice, then inhibited CXC chemokine receptor 3 (CXCR3) in ICI-treated mice to assess therapeutic effect on myocarditis phenotype. Furthermore, we delineated functional effects of CXCR3 blockade on T-cell and macrophage interactions in a transwell assay. We then correlated the results in human single-cell multi-omics data from blood and heart biopsy data from patients with ICI myocarditis.

**Results:** Single-cell multi-omics demonstrated expansion of CXCL9/10+CCR2+ macrophages and CXCR3hi CD8+ effector T-lymphocytes in the hearts of *MRL/Pdcd1-/-* mice correlating with onset of myocarditis development. Both depletion of CXCL9/10+CCR2+ macrophages and CXCR3 blockade respectively led to decreased CXCR3hiCD8+ T-cell infiltration into the heart and significantly improved survival. A transwell assay showed that selective blockade of CXCR3 and its ligand, CXCL10 decreased CD8+ T-cell migration towards macrophages, implicating this interaction in T-cell cardiotropism towards cardiac macrophages. Cardiac biopsies from patients with confirmed ICI myocarditis demonstrated infiltrating CXCR3+ lymphocytes and CXCL9+/CXCL10+ macrophages. Both mouse cardiac immune cells and patient peripheral blood immune cells revealed expanded TCRs correlating with CXCR3hi CD8+ T-cells in ICI myocarditis samples.

**Conclusions:** These findings bring forth the CXCR3-CXCL9/10 axis as an attractive therapeutic target for ICI myocarditis treatment, and more broadly, as a druggable pathway in cardiac inflammation.

## Introduction

Immune checkpoints are regulatory proteins on the surface of T-cells that prevent aberrant T-cell activation by binding to ligands on other cells, including tumor and antigen-presenting cells. Immunotherapy agents targeting these molecules, called immune checkpoint inhibitors (ICIs), are monoclonal antibodies that represent a major advancement in cancer treatment. ICIs block the interaction between immune checkpoints such as programmed cell death-1 (PD-1) and cytotoxic T-lymphocyte associated protein 4 (CTLA-4) and its ligands, thereby activating the immune system to kill tumor cells^1^. ICIs include pembrolizumab (anti-PD1), nivolumab (anti-PD-1), ipilimumab (anti-CTLA4), combination therapies, among others. The usage of ICIs to treat malignancies has increased from 13.2% to 28.2% of all oncologic therapies between 2017 and 2021^2^.

Unfortunately, the activation of T-cell immunity caused by ICIs can lead to immune related adverse events (irAEs) that affect many organs, including thyroid, lung, liver, colon, skin, and heart. Cardiac irAEs include pericarditis, vasculitis, myocardial infarction, and arrhythmias—but among the deadliest is myocarditis^1^. ICI myocarditis, although uncommon with a reported incidence of 0.04%-1.4%^3,4^, has an extremely high case fatality rate of up to 40-50%^5^ and can lead to life-threatening arrhythmias and heart failure. Research into the pathophysiology may identify strategies for earlier intervention and more effective treatments, preventing morbidity and mortality.

In general, myocarditis is an inflammatory disease affecting the myocardium, and despite its high mortality rate^6^, there are no current targeted therapies. It can result from several etiologies (i.e infectious pathogens, autoimmune disorders, drugs, etc.)^7,8^ and progress to dilated cardiomyopathy, heart failure, and even sudden death^9^. Macrophages and various T-cell subsets have been observed in the inflammatory response in autoimmune myocarditis^10^. Although steps have been made into understanding the pathophysiology of the disease, the underlying mechanisms behind myocarditis are still not fully known, and this prevents diagnosis and treatment, leading to poor patient prognosis. In identifying key molecular interactions, it will be possible to develop new targeted therapies for treatment.

Patients with ICI myocarditis have significant CD8+ cytotoxic T-cell and macrophage infiltrates in the myocardium^11,12^, indicating a possible role for both immune cell populations in the pathogenesis of the disease. In previously published work from our group^13^, analysis of blood from patients with ICI and blood/cardiac immune cells from a genetic knockout PD-1 knockout mouse model (*MRL/Pdcd1-/-*) using novel single-cell multi-omics technologies led to the discovery of a unique population of cytotoxic CD8+ T-cells re-expressing CD45RA (Temra) in patient samples and the correlating population of clonal effector CD8+ T-cells in the mouse model associated with the disease. We and others have observed that CD8+ T-cells and macrophages are significantly increased in heart biopsies taken from patients with ICI myocarditis as well the hearts of *MRL/Pdcd1-/-* mice, *Pdcd1-/-Ctla4+/-* mice, and A/J mice injected with PD-1 neutralizing monoclonal antibody, three established mouse models of ICI myocarditis^12–14^. Additionally, Ma et al. showed that the expansion of an inflammatory population of IFN-U-induced macrophages plays an important role in the pathogenesis of ICI myocarditis^15^.

Single-cell multi-omics technologies, such as single-cell RNA-sequencing (scRNAseq), single-cell T-cell receptor sequencing (scTCRseq), and indexing of transcriptomes and epitopes by sequencing (CITE-seq), can advance cell-specific understanding of the immunologic landscape of ICI-mediated myocarditis and elucidate key molecular mechanisms necessary for targeted therapeutics and precision medicine^16^. Our group specializes in high throughput single-cell multi-omics. While single-cell technologies can sometimes be limited by high cost, time-intensive work, and informatic requirements, we have found several ways to bypass these issues. Using methods such as a CITE-seq barcoded antibody multiplexing technique, we are able to combine multiple samples per lane/gel bead in the 10X Genomics single-cell RNA-seq pipeline^13^ and perform sequencing at a much higher throughput for greater statistical power. Here, we took advantage of the integration of high-throughput scRNAseq, scTCR-seq, and CITE-seq techniques to uncover and map specific cell populations and cell-cell interactions relevant to the pathogenesis of ICI myocarditis at a single-cell resolution.

We discovered a novel and critical T-cell/macrophage interaction downstream of IFN-U-induced macrophages, bringing forth a new targeted therapeutic option for ICI myocarditis treatment. We present the first extensive characterization of the interactions between CXCL9/10+CCR2+ macrophages and a subset of effector CD8+ T-lymphocytes that expresses CXC chemokine receptor 3 (CXCR3), potentiating ICI myocarditis. Interplay between chemokine receptors (i.e CXCR3) and their ligands (i.e. CXCL9 and CXCL10) are critical in cell migration and chemotaxis during immune cell trafficking, surveillance, and inflammation^17^. These findings shed new light on T-cell and macrophage interactions that propagate cardiac inflammation in this disease and advance understanding in the search for novel therapeutic targets.

We developed a novel pharmacological mouse model of anti-PD1 and anti-CTLA4 monoclonal antibody treatment that more accurately recapitulates the patient phenotype of ICI myocarditis. Using this mouse model and the murine model of PD-1 deletion on an MRL background (*MRL/Pdcd1-/-*)^13^, we identify the CXCR3-CXCL9/10 axis as a novel therapeutic target for treatment and further correlated these discoveries with peripheral blood mononuclear cells (PBMCs) and heart biopsy samples from patients with ICI myocarditis.

The interactions we uncover between chemokines and their receptors in T-cell and macrophage crosstalk are not only important for the search of a targeted therapeutic option for ICI myocarditis treatment but may also provide insight into the mechanisms of other cardio-immunological diseases.

## Results

### CXCL9/10+CCR2+ macrophages and clonally expanded CXCR3hiCD8+ T-cells are enriched in a mouse model of ICI myocarditis

To study immune checkpoint inhibitor myocarditis, we used a mouse model of PD-1 deletion on a Murphy Roths Large (MRL) background which develops fulminant myocarditis around four weeks of age with 70% penetrance (*MRL/Pdcd1-/-* mice). We performed single-cell multi-omics^18,19^ including scRNAseq, scTCR-seq, and CITE-seq on immune cells isolated from the heart and blood of the *MRL/Pdcd1-/-* and control MRL mice at time points 1, 2, and 4 weeks of age (Fig 1A). Unsupervised clustering of the sequential timepoint scRNAseq data revealed all major immune cell subpopulations (Fig 1B), which were identified using canonical markers. Feature plots showed robust expression of these markers (Fig1C, Supp Table 1A). Quantification of immune cell subpopulations also showed changes in T-cells, dendritic cells, and basophils at 2 weeks (Supp 1A).

**Figure 1.**
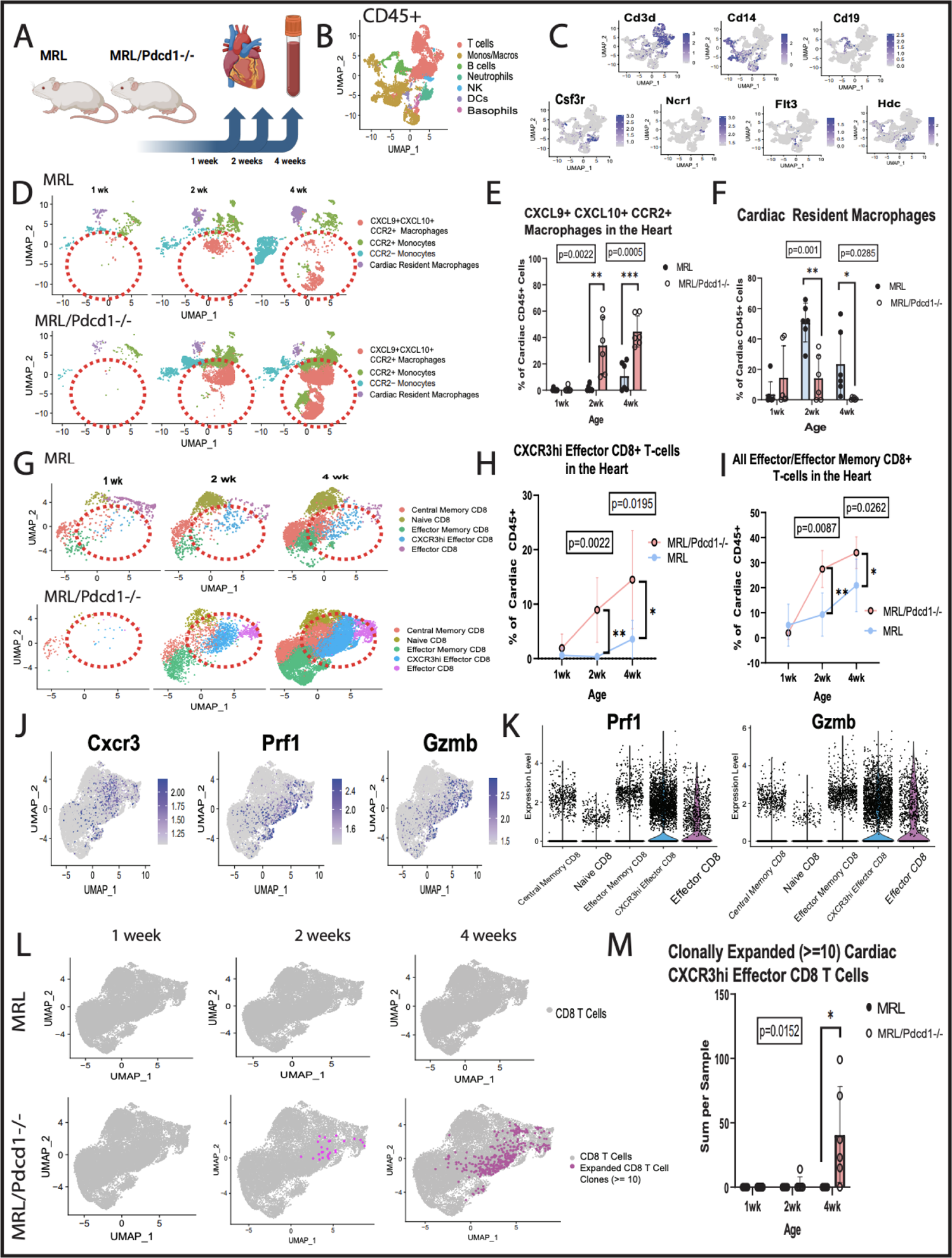
Identification of novel populations of CXCL9/10+CCR2+ macrophages and CXCR3hi CD8+ effector T-cells from single-cell sequencing data from MRL and *MRL/Pdcd1-/-* mice at timepoints 1 week, 2 weeks, and 4 weeks. A) Workflow schematic of time lapse experiment. Peripheral blood and hearts from MRL and *MRL/Pdcd1-/-* mice are harvested at ages 1 week (MRL heart: n=6, *MRL/Pdcd1-/-* heart: n=7, MRL blood: n=8, *MRL/Pdcd1-/-* blood: n=6), 2 weeks (MRL heart: n=6, *MRL/Pdcd1-/-* heart: n=6, MRL blood: n=6, *MRL/Pdcd1-/-* blood: n=6), and 4 weeks of age (MRL heart: n=6, *MRL/Pdcd1-/-* heart: n=6, MRL blood: n=6, *MRL/Pdcd1-/-* blood: n=6). **B)** UMAP of immune cell populations at the CD45+ level from scRNAseq of the heart and blood of MRL and *MRL/Pdcd1-/-* mice at ages 1, 2, and 4 weeks. **C)** Feature plots of immune cell markers at the CD45+ level. **D)** UMAP shows changes of macrophage subpopulations in the hearts and blood of MRL and *MRL/Pdcd1-/-* mice from age 1 week to 2 weeks to 4 weeks. **E)** Quantification of CXCL9/10+CCR2+ macrophages shows a significant increase in the hearts of *MRL/Pdcd1-/-* mice compared to MRL mice at 2 weeks (p=0.0022 by Mann-Whitney test) and 4 weeks (p=0.0005 by unpaired t-test). **F)** Quantification of cardiac resident macrophages shows a significant decrease in the hearts of *MRL/Pdcd1-/-* mice compared to that of MRL mice at 2 weeks (p=0.001 by unpaired t-test) and 4 weeks (p=0.0285 by unpaired t-test). **G)** UMAP shows changes of CD8+ T Cell subpopulations in the hearts and blood of MRL and *MRL/Pdcd1-/-* mice from age 1 week to 2 weeks to 4 weeks. **H)** Quantification of CXCR3hi CD8+ effector T-cells shows a significant increase in the hearts of *MRL/Pdcd1-/-* mice compared to that of MRL mice at 2 weeks (p=0.0022 by Mann-Whitney test) and 4 weeks (p=0.0195 by unpaired t-test). **I)** Quantification of all effector/effector memory CD8+ T-cells shows a significant increase in the hearts of *MRL/Pdcd1-/-* mice compared to that of MRL mice at 2 weeks (p=0.0087 by Mann-Whitney test) and 4 weeks (p=0.0262 by unpaired t-test). **J)** Feature plots indicate that CXCR3hi CD8+ effector T-cell populations strongly express cytotoxic markers. **K)** Violin plots of perforin and granzyme B show that expression is highest in CXCR3hi effector CD8+ and effector CD8+ subsets. **L)** UMAP at the CD8+ T-cell level shows expanded (>=10) cardiac CD8+ T-cell clones increase from 1 weeks to 4 weeks of age in MRL and *MRL/Pdcd1-/-* mice. **M)** Quantification of clonally expanded (>=10) cardiac CXCR3hi effector CD8+ T Cells shows a significant increase (p=0.0152 by Mann-Whitney test) in 4-week-old *MRL/Pdcd1-/-* mice. Biorender was used for panel **A.**

We then subdivided the monocyte/macrophage subset and performed unsupervised clustering to identify the monocyte and macrophage subpopulations based on canonical markers (Fig 1D, Supp 1B, Supp Table 1B). We observed that a population of CXCL9/10+CCR2+ macrophages was significantly increased in the hearts of *MRL/Pdcd1-/-* mice compared to the controls starting at two weeks of age and persisting to four weeks of age (Fig 1E). This correlates with the initiation of the myocarditis phenotype at 2 weeks of age in the *MRL/Pdcd1-/-* mice (Fig 1E). In contrast, the frequency of reparative LYVE1+ cardiac resident macrophages^20^ was significantly decreased in the hearts of *MRL/Pdcd1-/-* mice at timepoints 2 weeks and 4 weeks when compared to the control MRL (Fig 1F). We also noted that CCR2+ monocytes in the blood were significantly elevated in *MRL/Pdcd1-/-* mice at 4 weeks of age while the CCR2-monocytes were significantly decreased (Supp 1C).

To better understand the changes in CD8+ T-cell populations, we subdivided the T-cell cluster and performed unsupervised clustering to identify CD4+ and CD8+ subpopulations (Supp 1D) based off of canonical markers (Supp 1E, Supp Table 1C) The CD4+ T-cells remained unchanged, and the CD8+ T-cells were enriched in the *MRL/Pdcd1-/-* mice hearts at 2 weeks (Supp 1F). The CD8+ T-cell was further subdivided and annotated via canonical markers (Supp 1G, Supp Table 1D).

Unsupervised clustering revealed the emergence of a CXCR3hi CD8+ T-cell population in *MRL/Pdcd1-/-* mice hearts at 2 weeks, correlating with the expansion of the CXCL9/10 macrophage cluster (Fig 1G-H) and onset of myocarditis phenotype. The increased cardiac CD8+ T-cells were effector and effector memory cells, while naïve and central memory did not significantly increase (Fig 1I, Supp 1H). These effector CXCR3hi CD8+ T-cells showed high expression of cytotoxic markers such as *Prf1* and *Gzmb* (Fig 1J-K).

T-cell clonal expansion has been shown to play a role in autoimmunity, and expansion of select TCRs can indicate antigen specificity at a target site^21^. Single-cell TCR-sequencing revealed no baseline expansion of TCR sequences (defined here as greater than or equal to 10 clones) associated with CD8+ T-cells in the hearts of MRL mice, while expanded CD8+ T-cell clones begin to appear at 2 weeks of age and greatly proliferate by 4 weeks of age in the hearts of *MRL/Pdcd1-/-* mice as compared to control mice (Fig 1L-M).

### CXCR3hi CD8+ effector T-cells and CXCL9+CXCL10+ macrophages are predicted to interact in ICI myocarditis mice

After identification of novel CXCL9/10+CCR2+ macrophage and CXCR3hi effector CD8+ T-cell populations in the hearts of our *MRL/Pdcd1-/-* mice, we next utilized the bioinformatic package CellChatDB^22^ to further infer cell-cell communication and to predict signaling ligand-receptor interactions at different time points.

Analysis using CellChatDB revealed a predicted interaction between the CXCL9/10+CCR2+ macrophages in the heart and the CXCR3hi effector CD8+ T-cells in the blood that persists as the *MRL/Pdcd1-/-* increase in age from 1 to 2 to 4 weeks (Supp 2A). We found that the CXCL signaling pathway network showed high probability of communication between the CXCL9/10+CCR2+ macrophages and the CXCR3hi CD8+ effector T-cells at four weeks of age, and the CXCL9/10+CCR2+ macrophages were the source of the signal (Supp 2B). Furthermore, the communication probability between the CXCL9/10+CCR2+ macrophages and the CXCR3hi Effector CD8+ T-cells is higher in the *MRL/Pdcd1-/-* mice compared to MRL (Supp 2B). Then, focusing specifically on the CXCL9-CXCR3 and CXCL10-CXCR3 interactions in the CXCL signaling pathway in our four-week old mice, we observed that this signaling relationship is present between the CXCL9/10+CCR2+ macrophages and the CXCR3hi effector CD8+ T-cells in our *MRL/Pdcd1-/-* mice and absent in the MRL mice (Supp 2C).

### Macrophage depletion reduces cardiac immune cell infiltration in mice

In order to interrogate the role that macrophages play in ICI myocarditis, we treated *MRL/Pdcd1-/-* mice with liposomal clodronate to deplete macrophages compared to a liposomal control for two weeks (Fig 2A). Strikingly, we observed that mice treated with liposomal clodronate had a significantly higher probability of myocarditis-free survival compared to the controls (Fig 2B). Additionally, histological analysis showed that *MRL/Pdcd1-/-* mice treated with liposomal clodronate had significantly decreased overall immune cell infiltration compared to controls (Fig 2C, 2D).

**Figure 2.**
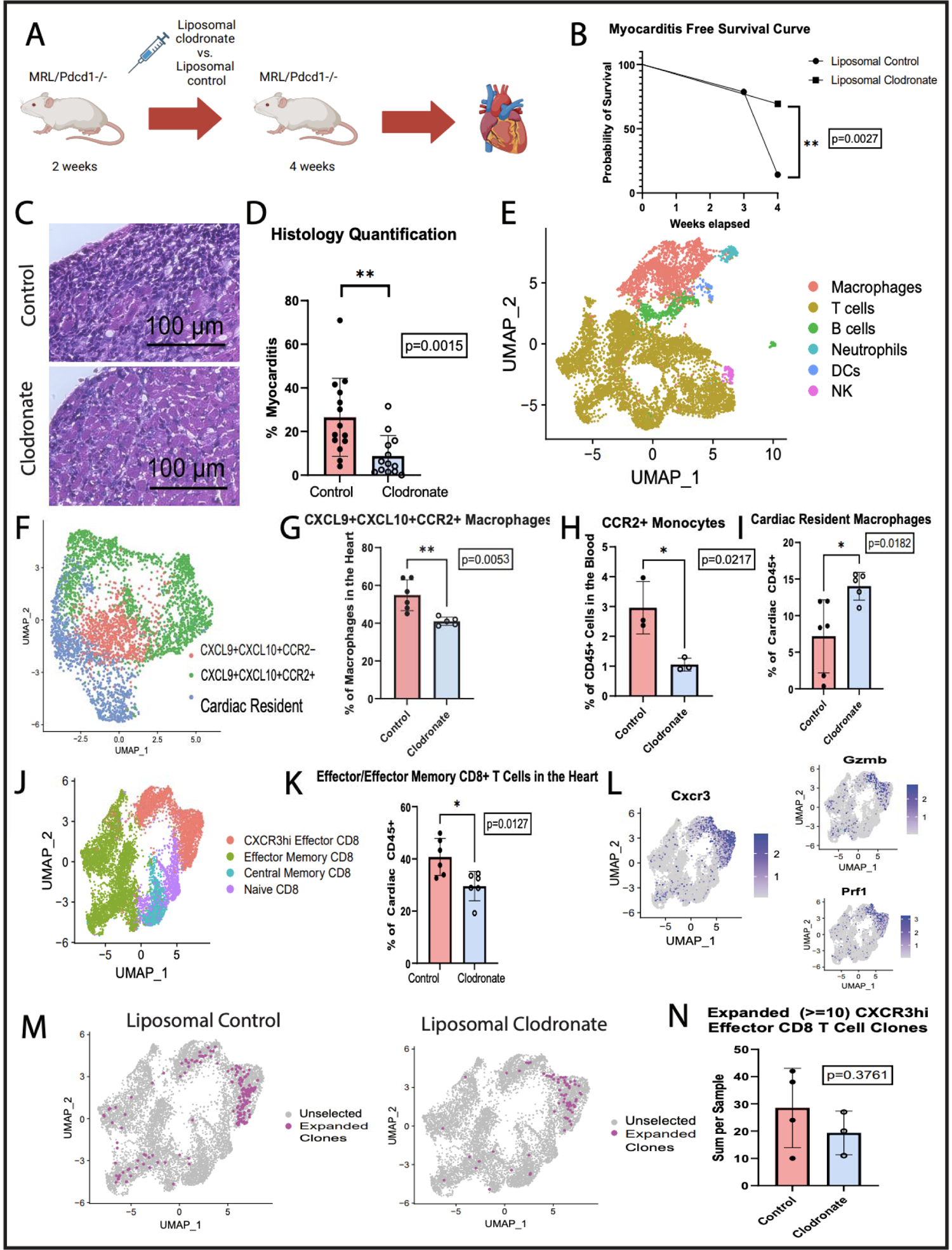
Depletion of monocytes/macrophages via liposomal clodronate attenuates ICI myocarditis in *MRL/Pdcd1-/-* mice. **A)** Workflow schematic of monocyte/macrophage depletion. *MRL/Pdcd1-/-* mice are injected every other day with liposomal control (n=6) or liposomal clodronate (n=6) to deplete monocytes/macrophages starting at 2 weeks of age. Hearts from these mice are harvested at 4 weeks of age. **B)** Myocarditis-free (<10%) survival curve of *MRL/Pdcd1-/-* mice injected with either liposomal control or liposomal clodronate. Mice injected with liposomal clodronate have a significantly higher probability of myocarditis-free survival (p=0.0027 by Log-rank test). **C)** Hematoxylin and eosin stain of hearts of *MRL/Pdcd1-/-* mice injected with either liposomal control or liposomal clodronate. **D)** Quantification of percentage of myocarditis in the hearts of *MRL/Pdcd1-/-* mice injected with either liposomal control or liposomal clodronate. Mice injected with liposomal clodronate have significantly less immune cell infiltration per area (p=0.0015 by Mann-Whitney test). **E)** UMAP of major immune cell populations identified at the CD45+ level from single-cell RNA-sequencing of the hearts of *MRL/Pdcd1-/-* mice injected with liposomal control or liposomal clodronate. **F)** UMAP of macrophage immune cell subpopulations in the hearts of *MRL/Pdcd1-/-* mice injected with liposomal control or liposomal clodronate. **G)** Quantification of CXCL9+CXCL10+CCR2+ macrophages analyzed via scRNAseq shows a significant decrease in the hearts of mice injected with liposomal clodronate compared to control (p=0.0053 by unpaired t-test). **H)** Quantification of CCR2+ monocytes analyzed via flow cytometry shows a significant decrease in the blood of mice injected with liposomal clodronate compared to control (p=0,0217 by unpaired t-test). **I)** Quantification of cardiac resident macrophages analyzed via scRNAseq shows a significant increase in the hearts of mice injected with liposomal clodronate compared to control (p=0.0182 by unpaired t-test). **J)** UMAP of CD8+ T-cell subpopulations in the hearts of *MRL/Pdcd1-/-* mice injected with liposomal control or liposomal clodronate. **K)** Quantification of all Effector and Effector Memory CD8+ T-cells in the hearts of *MRL/Pdcd1-/-* mice shows a significant decrease in that of those injected with liposomal clodronate compared to the control (p=0.0127 by unpaired t-test). **L)** Feature plots indicate that CXCR3hi CD8+ Effector T-cell populations strongly express cytotoxic markers such as Perforin and Granzyme B. **M)** UMAP at the CD8+ T-cell level shows expanded (>=10) cardiac CD8+ T-cell clones in *MRL/Pdcd1-/-* mice injected with either liposomal control or liposomal clodronate. Expanded TCRs are primarily concentrated in the CXCR3hi Effector CD8+ cluster. **N)** Quantification of clonally expanded (>=10) cardiac CXCR3hi Effector CD8+ T-cells shows a decreasing trend in *MRL/Pdcd1-/-* mice injected with liposomal clodronate compared with the control (p=0.3761 by unpaired t-test). Biorender was used for panel **A.**

We performed single-cell multi-omics including scRNAseq, scTCR-seq, and CITE-seq on immune cells isolated from the hearts of *MRL/Pdcd1-/-* mice treated with liposomal clodronate (n=6) or with control liposomes (n=6). Unsupervised clustering of the scRNAseq data revealed the major immune cell subpopulations (Fig 2E) which showed high expression of canonical markers (Supp 3A, Supp Table 2A). We found no significant changes in broad immune cell populations in the hearts of mice treated with liposomal clodronate versus control (Supp 3B). However, as presented below, we saw a decrease in CCR2+ monocytes in the blood and CXCL9/10+CCR2+ macrophage in the heart.

We subdivided the macrophages and performed unsupervised clustering to analyze the cardiac resident, CXCL9/10+CCR2+, and CXCL9/10+CCR2-macrophage subpopulations (Fig 2F, Supp 3C, Supp Table 2B). Quantification of the immune cell subsets revealed a significant decrease in the CXCL9/10+CCR2+ macrophage subpopulation in the hearts of *MRL/Pdcd1-/-* mice treated with liposomal clodronate (Fig 2G). We further analyzed circulating macrophage/monocytes by flow cytometry (Supp 3D), revealing that CCR2+ monocytes were significantly decreased in the blood of mice treated with clodronate (Fig 2H). We also see a corresponding significant increase in relative frequency of anti-inflammatory TIMD4+ cardiac resident macrophages in the hearts of mice treated with liposomal clodronate (Fig 2I). Quantification of other macrophage subpopulations had no significant change (Supp 3E).

We then sought to test if the decrease in pro-inflammatory CXCL9/10+CCR2+ macrophages had an effect on the CD8+ T-cell subpopulations we previously identified in the heart. We subdivided the T-cell population and annotated CD4+ and CD8+ after unsupervised clustering of the immune cells (Supp 3F, Supp Table 2C, Supp 3G).

Quantification of bulk CD3+ T-cells did not show significant changes (Supp 3H), however we found upon unsupervised clustering of CD8+ T-cells that effector and effector memory CD8+ T-cells were significantly decreased in the hearts of *MRL/Pdcd1-/-* mice treated with liposomal clodronate (Fig 2J-2K, Supp 3I, Supp Table 2D). Further, the decreased CXCR3hi CD8+ effector T-cell population showed high expression of cytotoxic markers perforin and granzyme B (Fig 2L). Quantification of other CD8+ T-cell subpopulations showed no significant changes (Supp 3J).

Single-cell TCR-sequencing similarly revealed that expanded T-cell clones overlap with the labeled CXCR3hi CD8+ effector T-cells (Fig 2M). Mice treated with liposomal clodronate showed a trend in decreased clonal expansion CD8+ T-cells in their hearts compared to the control mice (Fig 2N).

### *In Vivo* CXCR3 Blockade Attenuates the Myocarditis Phenotype

To complement our genetic mouse model for ICI myocarditis, we established a new pharmacological mouse model by injecting MRL mice with monoclonal antibodies against PD1 and CTLA-4 twice a week (Supp 4A), which better models the combination immunotherapy that is a significant risk factor for ICI myocarditis in patients. We found that mice develop myocarditis after 6 doses of combination immunotherapy with an incidence of 25-50%, and flow cytometry (Supp 4B) indicated an increasing trend in CD8+ T-cells and macrophages in the hearts of mice treated with combination immunotherapy (Supp 4C). Histological analysis likewise shows infiltration of immune cells and significant myocardial damage in the hearts of these immunotherapy-treated mice (Supp 4D).

We then sought to test if inhibition of the CXCL9/10-CXCR3 axis we have characterized thus far affected the myocarditis phenotype. We utilized a CXCR3 blockade by injecting MRL mice with combination immunotherapy as well as with either an anti-CXCR3 monoclonal antibody or an IgG control (Fig 3A) to inhibit the interaction between CXCR3-positive T-cells with its ligands.

**Figure 3.**
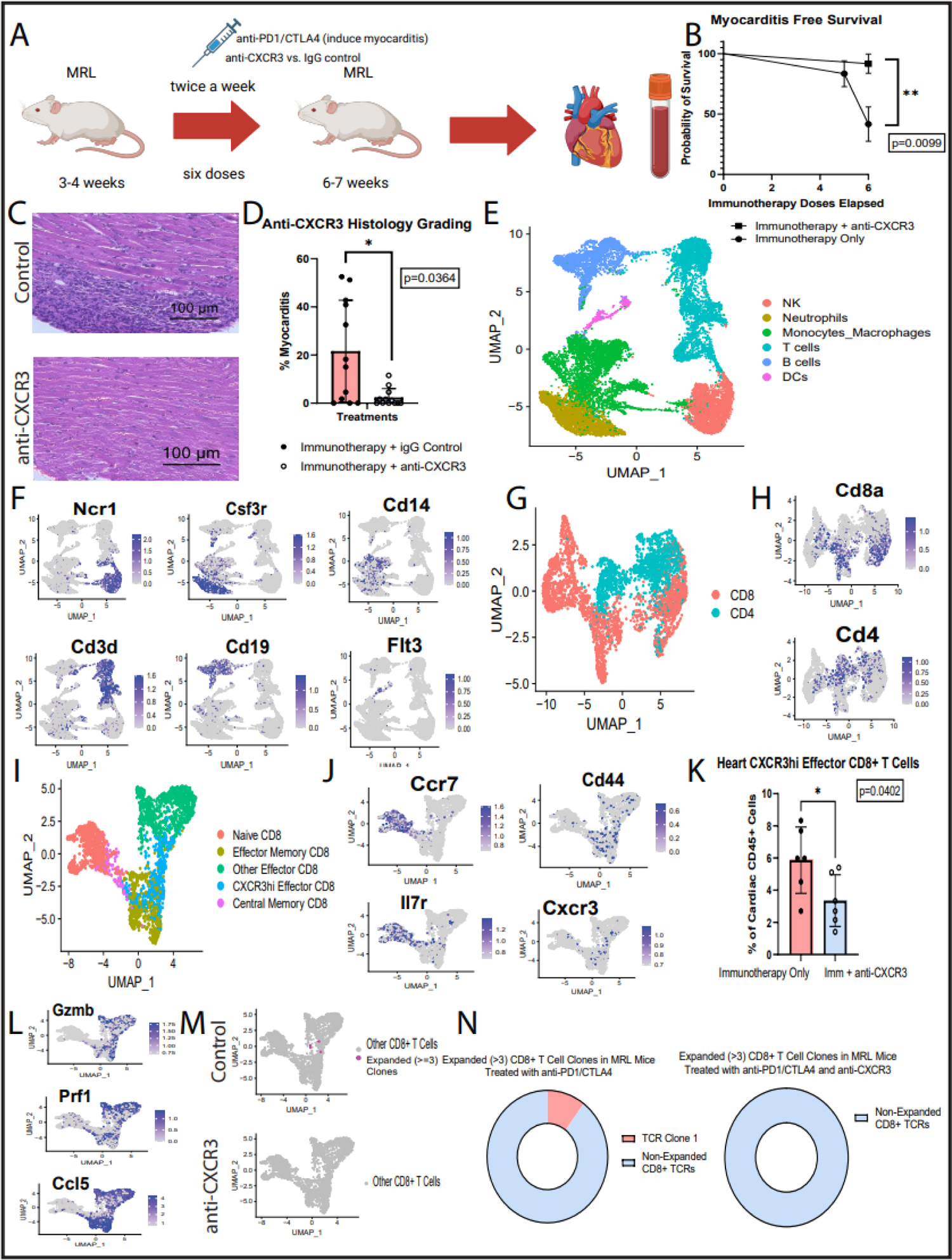
*In vivo* CXCR3 blockade in a pharmacological mouse model for ICI myocarditis significantly decreases immune cell infiltration in the heart. A) Workflow schematic of *in vivo* CXCR3 blockade in MRL mice treated with anti-PD1 and anti-CTLA4 combination immunotherapy. MRL mice are treated with immunotherapy and either anti-CXCR3 (n=6) or an IgG control (n=6). Heart and blood are harvested after six doses of immunotherapy. **B)** Myocarditis-free (<10%) survival curve of immunotherapy-treated MRL mice injected with either anti-CXCR3 or IgG control. Mice injected with anti-CXCR3 have a significantly higher probability of myocarditis-free survival (p=0.0099 by Log-rank test). **C)** Hematoxylin and eosin staining of hearts of immunotherapy-treated MRL mice injected with either anti-CXCR3 or IgG control. **D)** Quantification of percentage of myocarditis in the hearts of immunotherapy-treated MRL mice injected with either anti-CXCR3 or IgG control. Mice injected with anti-CXCR3 have significantly less immune cell infiltration per area (p=0.0364 by Mann-Whitney test). **E)** UMAP of major immune cell populations identified at the CD45+ level from single-cell RNA-sequencing of the hearts and blood of immunotherapy-treated MRL mice injected with either anti-CXCR3 (n=6) or IgG control (n=6) as well as the hearts of MRL mice treated with only IgG control (n=3). **F)** Feature plots of major immune cell markers for the labeled populations at the CD45+ level. **G)** UMAP of T-cell subpopulations at the CD3+ level. **H)** Feature plots of CD4+ and CD8+ markers at the CD3+ level. **I)** UMAP of CD8+ T Cell subpopulations in the hearts and blood of immunotherapy-treated MRL mice injected with either anti-CXCR3 or IgG control. **J)** Feature plots of key markers for CD8+ T-cell subpopulation annotations. **K)** Quantification of CXCR3hi Effector CD8+ T-cells shows a significant decrease in the hearts of mice injected with anti-CXCR3 (p=0.0402 by unpaired t-test). **L)** Feature plots indicate that effector and effector memory CD8+ T-cell subsets express cytotoxic markers Granzyme B and Perforin, as well as chemotactic markers such as Ccl5. **M)** UMAP at the CD8+ T Cell level shows expanded (>3) cardiac CD8+ T-cell clones in immunotherapy-treated MRL mice injected with either anti-CXCR3 or IgG control. There is decreased expansion in mice treated with anti-CXCR3. Expanded TCRs are primarily concentrated in the CXCR3hi Effector CD8+ cluster. **N)** Pie chart of expanded (>3) CD8+ T-cell Clones in immunotherapy-treated MRL mice injected with either anti-CXCR3 or IgG control. Biorender was used for panel **A.**

We found that mice treated with combination immunotherapy and anti-CXCR3 had a significantly higher probability of myocarditis-free survival when compared to mice that were treated only with immunotherapy (Fig 3B). We found a significant decrease in immune cell infiltration in the hearts of these (Fig 3C). Histologic scoring of the myocardium revealed a significant decrease in myocarditis in mice treated with the CXCR3 blockade (Fig 3D).

We performed single-cell multi-omics including scRNAseq, scTCR-seq, and CITE-seq on immune cells isolated from the hearts and blood of MRL mice treated with combination immunotherapy and anti-Cxcr3 (n=6) or IgG control (n=6) as well as on mice treated only with IgG (n=3). Unsupervised clustering of the isolated immune cells revealed all major immune cell subpopulations were present (Fig 3E, Supp Table 3A). Annotations of the immune cell subpopulations were based off high expression of canonical cell markers (Fig 1F). There were no significant changes in immune cell frequency at the CD45+ level (Supp 4E).

We then subdivided the T-cell population and performed unsupervised clustering on the subset to label CD4+ versus CD8+ T-cells (Fig 3G) via RNA expression of canonical markers (Fig 3H, Supp Table 3B). Quantification of these subpopulations can be found in the supplementary figures (Supp 4F). The CD8+ T-cells were subdivided and unsupervised clustering performed to identify CD8+ T-cell subpopulations (Fig 3I).

Naive, central memory, effector memory, CXCR3hi effector, and effector CD8+ T-cells were annotated based on canonical cell markers (Fig 3J, Supp Table 3C). CXCR3 blockade resulted in significantly lower frequencies of specifically CXCR3hi effector CD8+ T-cells in the heart (Fig 3K). The specificity of our CXCR3 blockade was highlighted by no significant changes to other CD8+ T-cell subpopulations or CD4+ T-cells in the heart with anti-CXCR3 treatment (Supp 4G). Additionally, CXCR3hi CD8+ T-cell frequency did not change in the blood (Supp 4H), indicating no depletion of CXCR3hi CD8+ T-cells with this monoclonal antibody.

The CXCR3hi CD8+ populations impacted by the antibody blockade were also positive for cytotoxic markers perforin and granzyme B, as well as chemotactic markers such as *Ccl5*, indicating our treatment specifically impaired the cardiotropism of highly cytotoxic T-cells (Fig 3L).

Single-cell TCR-sequencing revealed that clonally expanded (>3 TCRs) CD8+ T-cells in the heart overlap with the CXCR3hi CD8+ T-cells (Fig 3M). Importantly, the mice that were treated with CXCR3 blockade had decreased numbers of expanded CD8+ TCRs in the heart, confirming that blocking the CXCR3 axis decreased specifically the clonally expanded T-cells that traffic to the heart (Fig 3N).

### *In Vitro* Blockade of CXCR3 and CXCL10 Decreases Migration of CD8+ T-cells towards Macrophages

In order to determine the role of CXCR3, CXCL9, and CXCL10 in CD8+ T-cell chemotaxis towards macrophages, we utilized a transwell assay system. We isolated CD8+ T-cells from the heart and macrophages from *MRL/Pdcd1-/-* mice, plated CD8+ T-cells on the top insert and macrophages in the bottom well, then measured CD8+ T-cell migration towards the macrophages in the presence or absence of CXCR3 and CXCL9/10 blockade (Fig 4A).

**Figure 4.**
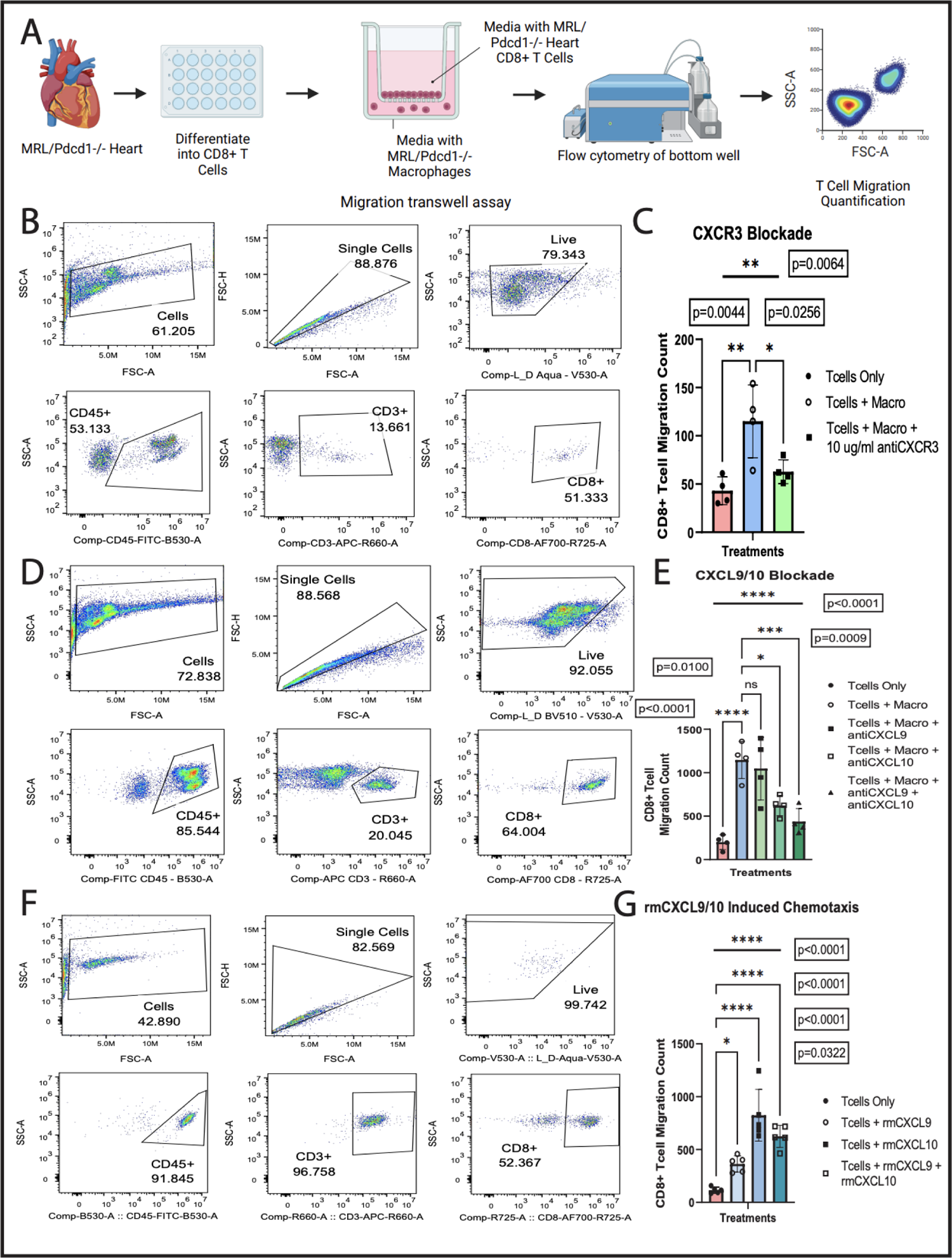
Selective blockade of CXCR3, CXCL9, and CXCL10 in a transwell system decreases CD8+ T-cell migration towards macrophages. **A)** Workflow schematic of *in vitro* transwell assay of T-cell and macrophage crosstalk. CD8+ T-cells are differentiated from the hearts of *MRL/Pdcd1-/-* mice and plated in the top insert of a transwell. Peritoneal macrophages from *MRL/Pdcd1-/-* mice are plated in the bottom well. Migration of CD8+ T-cells is assessed via flow cytometry of the bottom well. **B)** Flow cytometry gating of *in vitro* CXCR3 blockade of CD8+ T-cells. **C)** Quantification of CD8+ T-cell migration after CXCR3 blockade of the CD8+ T-cells in the upper well (p=0.0064 by ordinary one-way ANOVA). There is a significant increase in migration of CD8+ T-cells in the presence of macrophages (p=0.0044 and p=0.0256 by Dunnett’s multiple comparisons test), and this migration is significantly decreased when CD8 T-cells are incubated with anti-CXCR3 (p=0.0256 by Dunnett’s multiple comparisons test). **D)** Flow cytometry gating of *in vitro* CXCL9 and CXCL10 blockade of macrophages. **E)** Quantification of CD8+ T-cell migration after CXCL9 and/or CXCL10 blockade of the macrophages in the bottom well (p<0.0001 by ordinary one-way ANOVA). There is a significant increase in migration of CD8+ T-cells in the presence of macrophages (p<0.0001 by Dunnett’s multiple comparisons test). Blockade of CXCL9 has no significant change in migration (p=0.9051 by Dunnett’s multiple comparisons test). Blockade of CXCL10 results in significant decrease in CD8+ T-cell migration towards macrophages (p=0.0100 by Dunnett’s multiple comparisons test). Blockade of both CXCL9 and CXCL10 results in significant decrease in migration as well (p=0.0009 by Dunnett’s multiple comparisons test). **F)** Flow cytometry gating of *in vitro* rmCXCL9 and rmCXCL10 induced migration of CD8+ T-cells. **G)** Quantification of CD8+ T-cell migration in the presence of recombinant mouse protein CXCL9 and/or CXCL10 (p<0.0001 by ordinary one-way ANOVA). Addition of rmCXCL9 to the bottom well significantly increases migration of CD8+ T-cells to the bottom well (p=0.0322 by Dunnett’s multiple comparisons test). Addition of rmCXCL10 to the bottom well significantly increases migration of CD8+ T-cells as well (p<0.0001 by Dunnett’s multiple comparisons test), and addition of both ligands results in significant migration (p<0.0001 by Dunnett’s multiple comparisons test). Biorender was used for panel **A.**

We observed that there was a significant increase in migration of CD8+ T-cells towards the bottom well in the presence of macrophages (Fig 4B). However, blockade of the CXCR3 receptor on the CD8+ T-cells significantly decreased migration towards the macrophages (Fig 4B-4C).

Using selective blockade of CXCL9 and CXCL10, we determined that only blocking CXCL9 did not significantly impact CD8+ T-cell migration towards macrophages. However, inhibition of CXCL10 did significantly decrease CD8+ T-cell migration towards the macrophages, which was modestly amplified by combined inhibition of both CXCL9 and CXCL10 (Fig 4D-4E).

To further test the importance of CXCL9/10 in T-cell migration towards cardiac macrophages, we performed the same chemotaxis assay with the addition of recombinant CXCL9 (rmCXCL9) and CXCL10 (rmCXCL10). Both rmCXCL9 and rmCXCL10 induced a significant increase in CD8+ T-cell migration in comparison to a control containing only media in the bottom well. rmCXCL10 was observed to induce migration to a greater degree than rmCXCL9, with no significant additive effects of combining both chemokines (Fig 4F-4G).

### ICI Myocarditis Patient Heart Biopsies Stain Positive for CXCR3, CXCL9, and CXCL10

To investigate if the findings of CXCL9/10+ macrophages and CXCR3+CD8+ T-cells can be translated to patients with ICI myocarditis, we probed existing single-cell sequencing data from patients with and without ICI myocarditis previously published by our group^13^. The cohort included 30 patients, including healthy controls not on ICIs (n=6), Group A of patients treated with ICIs but having no irAEs (n=8), Group B of patients treated with ICIs and having non-myocarditis irAEs (n=8), and Group C of patients that were treated with ICIs and had ICI myocarditis (n=8) (Fig 5A).

**Figure 5.**
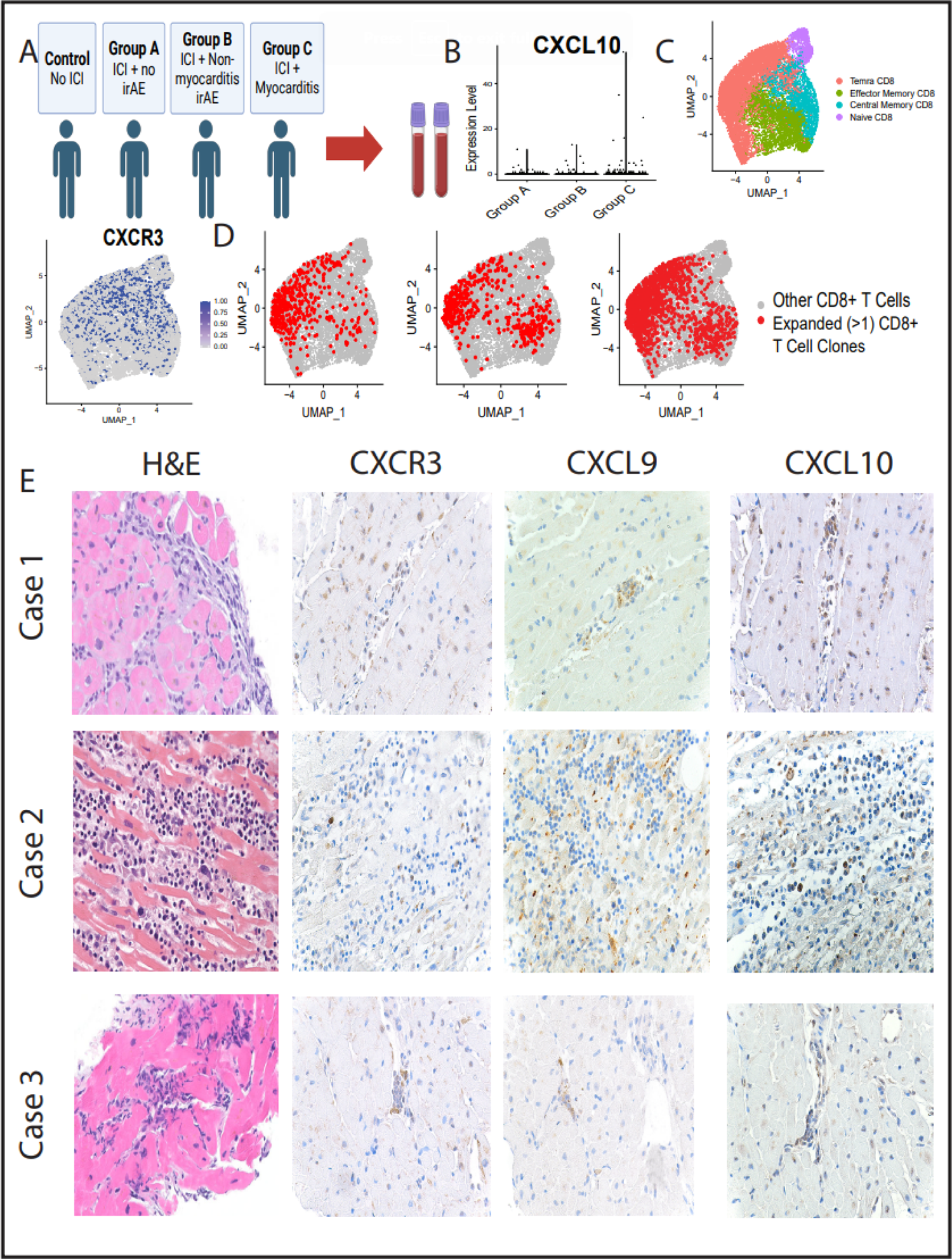
Heart and blood samples from patients with ICI myocarditis reveals CXCR3hi CD8+ T-cells and CXCL9/10+ macrophage populations. **A)** Single-cell sequencing data of human samples included PBMCs from healthy controls not on ICIs (n=6), Group A of patients treated with ICIs but having no irAEs (n=8), Group B of patients treated with ICIs and having non-myocarditis irAEs (n=8), and Group C of patients that were treated with ICIs and had ICI myocarditis (n=8). **B)** Violin plots of CXCL10 expression in the macrophages/monocytes of human PBMCs. **C)** Feature plot of CXCR3 expression on a UMAP of annotated CD8+ T-cell subpopulations shows an overlap with immune cells previously labeled as Temra CD8+. **D)** Single-cell TCR-sequencing of clonally expanded (>1) CD8+ T-cell clones in the different patient groups. **E)** Hematoxylin and eosin staining and immunoperoxidase staining for CXCR3, CXCL9, and CXCL10 in the hearts of three patients with confirmed ICI myocarditis. The lymphocytes show immunoreactivity for CXCR3 and the macrophages stained show CXCL9 and CXCL10 expression. Biorender was used for panel **A.**

We probed macrophages for CXCL10 expression and found an increase in expression in PBMCs from patients with ICI myocarditis (Group C) compared to patients without irAEs (Group A) and patients with non-myocarditis irAEs (Group B) (Fig 5B). Further, when we subdivided the T-cell compartment, we found a high proportion of CXCR3+CD8+ T effector cells re-expressing CD45RA (Temra CD8+), which were previously found to be significantly increased in Group C compared to healthy controls as well as Group A and B^13^ (Fig 5C). Additionally, single-cell TCR-sequencing analysis also showed that the CD8+ T-cell clones increased from Group A to Group B to Group C (Fig 5D) and overlap with CXCR3hi CD8+ T-cells (Fig 5C).

Finally, we examined three heart biopsies taken from patients with confirmed ICI myocarditis at the histologic level, and as expected found significant immune cell infiltration among cardiomyocytes. Further, we found that high numbers of CXCR3 positive lymphocytes as well as macrophages positive for CXCL9 and CXCL10 (Fig 5E), This data confirms that the prominent populations of CXCL9CXCL10+ macrophages and CXCR3hi CD8+ T-cells in the hearts of our ICI myocarditis mouse model are present in human samples as well.

## Discussion

In this study, we establish a role for CD8+ T-cell and macrophage crosstalk in the pathogenesis of immune checkpoint inhibitor-mediated myocarditis. Sequential time point single-cell multi-omics of cardiac immune cells from mice deficient in PD-1 revealed expansion of CXCR3hi CD8+ effector T-cells and CXCL9/10+CCR2+ macrophages in the heart as the disease progresses. Macrophage depletion resulted in significant mitigation in disease phenotype and increase in myocarditis-free survival, confirming CXCL9/10+CCR2+ macrophages as necessary in the progression of ICI myocarditis. Alongside these changes to macrophages, we experimentally determine for the first time that *in vivo* CXCR3 blockade mitigates ICI myocarditis and cardiac CD8+ CXCR3hi T-cell infiltration, bringing forth an important new therapeutic target. We also use *in vitro* CXCR3, CXCL9, and CXCL10 blockade to mechanistically confirm the importance of CXCR3-CXCL9/10 interactions in cardiac CD8+ T-cell migration towards macrophages. Lastly, we correlated the CXCR3hi CD8+ T-cells and CXCL9/10+ macrophages in our mice with corresponding cell populations in peripheral blood^13^ and in heart biopsies of patients with ICI myocarditis.

Our findings introduce the CXCR3-CXCL9/10 axis as a critical target in the pathophysiology of ICI myocarditis for precision medicine-based therapeutic treatment. It has been proposed that T-cells target a shared antigen between tumor and heart^11^. Once activated, these T-cells release IFN-U, and IFN-U-receptive CCR2+ macrophages differentiate into CXCL9/10+ macrophages^15^. We show that in our ICI myocarditis mouse model these CXCL9/10+CCR2+ macrophages and a population of CXCR3hi CD8+ effector T-cells are significantly elevated in the heart. These CXCR3+ effector CD8+ T-cells also express high levels of cytotoxic markers granzyme B and perforin^13^ and likely contribute to cardiac damage (Figure 6). By blocking the interaction between CXCR3+ cytotoxic CD8+ T-cells and CXCL9/10 released by macrophages, we successfully prevented and treated ICI myocarditis in our mouse model, thus implicating this axis as a vital drug target. This marks an important step in the field of ICI myocarditis management due to current limited evidence-based targeted therapies for the disease.

**Figure 6.**
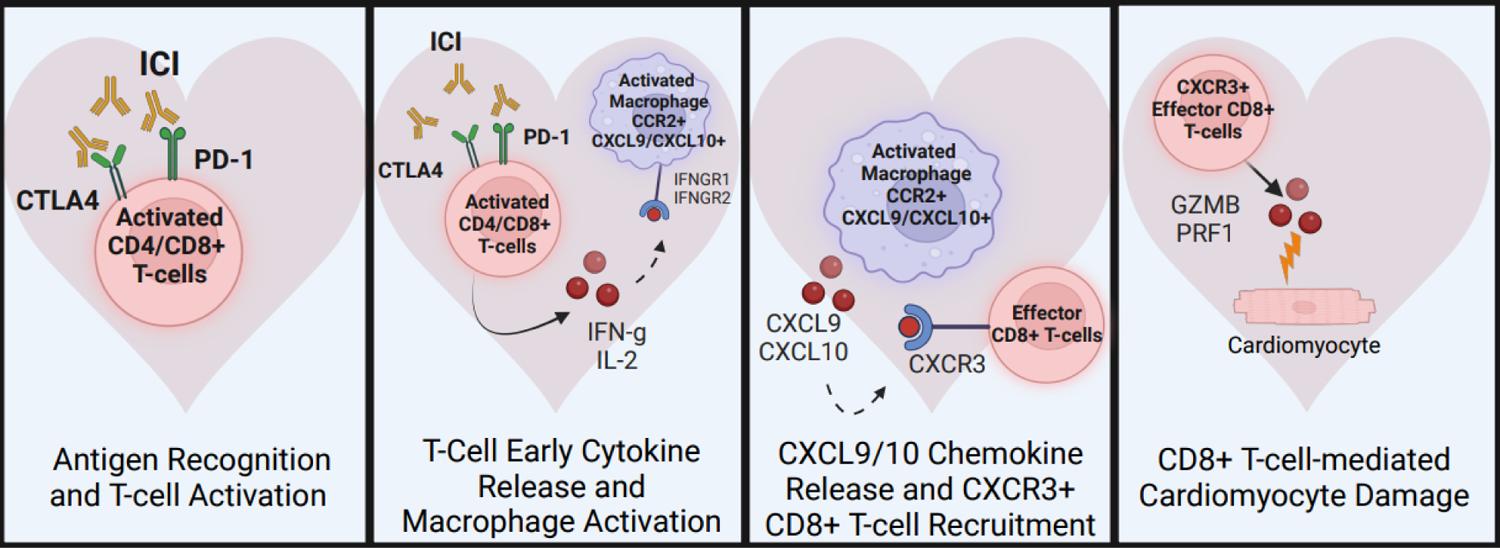
Diagram of the pathogenesis of ICI myocarditis. Biorender was used to create this figure.

The current first-line immunosuppressive therapy for ICI myocarditis is corticosteroids, and if this is not sufficient, then other immunosuppressants such as mycophenolate mofetil, abatacept^23^, and JAK inhibitors^1^ have been attempted. Abatacept is FDA approved for rheumatoid arthritis and is currently in clinical trials for ICI myocarditis^24,25^. JAK inhibitors, including ruxolitinib^26^ and tofacitinib^27^, have also shown promise for mitigating ICI myocarditis through observational studies. Alemtuzumab^28^, a monoclonal antibody for CD52, also has case study level data. However, there remains a need to find additional targeted therapeutic options with potential for temporal control.

Targeted therapy based on inhibiting chemokine receptors or specific chemokine ligand/receptor conformations can be used to control cellular responses for precision-based treatment. Because chemokines have a very short half-life *in vivo*, this makes CXCR3 and its ligands an attractive target for increased temporal control, an important consideration in cancer treatment. The ability for CXCR3 inhibition to maximize short-term therapeutic effects on the inflamed heart while minimizing long-term undesirable immunosuppressive effects in non-target tissue such as tumor may make it a well-tolerated therapy^29^. Further studies will help delineate the efficacy of chemokine receptor and chemokine therapies as a either a temporary short-term or even preventative method against ICI myocarditis.

For any immunosuppressive therapy in ICI myocarditis, important consideration must be given to its potential impact on tumor infiltrating leukocytes in disease control^30^. Although the role of chemokines/chemokine receptors in tumor biology is complicated, it is possible that blockade of CXCR3 and its ligands may actually play a positive role in tumor control. For example, CXCR3 inhibition improved NK cell-based adoptive immunotherapy for multiple myeloma^31^. Another study showed that CXCR3 blockade inhibited lung metastasis in a mouse model of metastatic breast cancer^32^, and it has been suggested that CXCR3 can be targeted to suppress lymph node metastasis of colon cancer^33^. It is also relevant to note that CXCL10 blockade effects may depend on the corresponding CXCR3 receptor isoform variant (CXCR3-A vs. CXCR3-B), necessitating further study^34^. In fact, CXCR3-A has the highest expression among the isoforms, and also the highest affinity for CXCL9/10/11. Simultaneously, CXCR3-A also facilitates proliferation and migration of tumor cells, making it a useful target for cancer therapy^35^. In general, it is especially important to understand the differing roles of CXCR3 and its ligands as they pertain to both tumor prognosis and ICI myocarditis outcome, but the two goals may not necessarily be conflicting if the correct target is chosen. Future studies will be needed to further elucidate the detailed responses of heart vs. tumor infiltrating lymphocytes to CXCR3 axis inhibition. Regardless, here we present the first important step towards investigating the therapeutic nature of CXCR3 axis inhibition in ICI myocarditis treatment.

Importantly, the CXCR3 axis may be a potential therapeutic target for not only ICI myocarditis but other inflammatory cardiac pathologies as well. In general, CXCL10 has been noted to be elevated in the heart in rodent models of myocarditis^36^. In SARS-CoV-2 mRNA vaccine-associated myocarditis, circulating CXCL10 was found to be elevated in the plasma^37^. In non-myocarditis heart disease, plaque progression in coronary artery disease has been evidenced to show high CXCR3 expression in immune cell buildup, and CXCL10 is reported to be elevated in circulation^36^. In mouse models of transverse aortic constriction for heart failure, increased circulating levels of CXCL9 and CXCL10 have also been observed^36^. Studies have shown that peripheral levels of CXCL10 were significantly higher in patients with stable and unstable angina in comparison to a healthy control group^38^. In nonischemic heart failure, it has been observed that cardiac myeloid cells and fibroblasts produced CXCL9 and CXCL10 in response to cardiac pressure overload. *Cxcr3-/-* transverse aortic constriction (TAC) mice were seen to have decreased CXCR3+CD4+ T-cell infiltration that was associated with cardiac dysfunction^39^. In patients with left ventricular dysfunction and heart failure, CXCL9 and CXCL10 were increased significantly in symptomatic patients^40^.

It is also noteworthy that CXCR3 antagonists have been proposed for use to treat inflammatory and autoimmune diseases and have been well-tolerated^41^. ACT-777991^42^ was tested in Type I diabetes patients to reduce migration of CXCR3-expressing cells to the pancreas. During the study, no adverse events of moderate or severe intensity occurred, and it was overall seen to be safe and well tolerated across the tested dose range. CXCL10 blockades have been in clinical trials for other diseases. MDX-1100^43^, a human monoclonal antibody for CXCL10, in combination with methotrexate has shown clinical efficacy in treating patients with rheumatoid arthritis. Another example is BMS-936557^44^, which was given to patients with ulcerative colitis in a double-blind, multicenter, randomized study, and the drug was correlated with histological improvement and increased clinical response. It appeared to be generally safe, and well-tolerated.

To conclude, we performed *in vivo* blockade of the CXCR3 axis with novel high-dimensional single-cell sequencing and scTCR-sequencing to map specific T-cell populations temporally and within the disease organ. We also performed *in vitro* transwell migration assays to confirm functional significance of the blockade on T-cell and macrophage crosstalk. Additionally, we confirmed our findings with translational patient data. These findings highlight CXCR3 axis blockade as an attractive novel therapeutic strategy in ICI myocarditis, with potential broad-ranging implications in other cardiac inflammatory and autoimmune diseases. In doing so, we lay paradigm-shifting groundwork for further studies in chemokine and chemokine receptor therapies in the heart and other organs affected by T-cell mediated inflammation.

## Novelty and Significance

### What is known?

- ICI myocarditis is associated with an increase in both cytotoxic CD8+ T-cells and macrophages in patients and preclinical models, suggesting a role for crosstalk between T-cells and macrophages potentiating cardiac inflammation.

### What new information does this article contribute?

- CXCL9/CXCL10+CCR2+ macrophages and CXCR3hiCD8+ T-cells are significantly increased in the hearts of mice with ICI myocarditis, and correspondingly, CXCL9/10+ macrophages and CXCR3+ lymphocytes are present in the hearts of patients with ICI myocarditis.
- Both macrophage depletion and CXCR3 blockade in a novel pharmacological mouse model of ICI myocarditis reduces myocarditis phenotype.
- Inhibition of both CXCR3 and its ligand CXCL10 respectively decreases migration of cardiac CD8+ T-cells towards macrophages.

Using single-cell multi-omics in both a genetic and novel pharmacological mouse model of ICI myocarditis, we uncover significantly increased populations of CXCL9/10+CCR2+ macrophages and CXCR3hiCD8+ effector T-cells in the hearts of mice with ICI myocarditis. We correlated this discovery with patient single-cell PBMC data as well as with patient heart biopsy samples. We show experimentally for the first time that depletion of CCR2+ macrophages and blockade of the chemokine receptor CXCR3 results in a significant reduction of ICI myocarditis phenotype and cardiac-infiltrating CXCR3hiCD8+ T-cells in mice. By elucidating downstream mechanisms of IFN-U-induced CXCL9/10+CCR2+ monocyte-derived macrophages, we discover the significance of the CXCR3-CXCL9/10 pathway in the progression of ICI myocarditis and bring forth CXCR3 and its ligands as a novel therapeutic target for treatment of the disease. The chemokine axis CXCR3-CXCL9/10 is an attractive therapeutic option for ICI myocarditis and, in general, for cardiac inflammatory diseases.

## Materials and Methods

### Data Availability

Flow cytometry data can be found on Mendeley Data at doi: 10.17632/8b7cy856X2.1. Single-cell RNA sequencing, single-cell TCR sequencing, and CITE-seq data can be found in the Gene Expression Omnibus under accession number GSE250056.

### Statistical Methods

Single-cell statistical analysis, visualization, and graph generation was performed in R using the Seurat^45,46^ package. Other statistical analyses were performed in GraphPad Prism 9.5.1. P value significance was categorized as nonsignificant (P >= 0.05), *P < 0.05, **P<0.01, ***P<0.001, and ****P<0.0001.

### Mice

We used Murphy Roths Large (MRL) mice as our control population and a genetic mouse model of PD-1 deletion on the MRL background (*MRL/Pdcd1-/-*)^13,47^ for our experimental myocarditis model. For our pharmacological model, MRL mice were injected intraperitoneally with 400 ug of anti-PD1 monoclonal antibody and 400 ug of anti-CTLA4 monoclonal antibody (BioXCell) or 800 ug of IgG control antibody twice a week.

### Macrophage Depletion

*MRL/Pdcd1-/-* mice were injected intraperitoneally with 100 ul of 5 mg/ml control PBS liposomes or liposomal clodronate^48,49^ (LIPOSOMA) every 2 days starting at 2 weeks of age to deplete macrophages.

### *In Vivo* CXCR3 Blockade in MRL Mice treated with anti-PD1 and anti-CTLA4

Anti-CXCR3 monoclonal antibody (BioXCell; Cat#: BE0249) or IgG control (BioXCell; Cat#: BE0091) was administered at a dosage of 200 ug, starting two days before immunotherapy treatment and then twice a week at the same time as anti-PD1/anti-CTLA4 treatment.

### *In Vitro* Transwell System for CXCR3, CXCL9, CXCL10 Blockade

Cardiac CD8+ T-cells^50^ were incubated with 10 ug/ml of anti-CXCR3 (BioXCell) or IgG control (BioXCell) for 30 minutes at 37C and added to the top insert of a 96 well transwell (Corning, 3387). Peritoneal macrophages^51^ from *MRL/Pdcd1-/-* mice were plated in the bottom well of the transwell. For the CXCL9 blockade, macrophages were incubated with 3 ug/ml of anti-CXCL9 (R & D; Cat #: AF492SP) for 45 minutes at 37C. For the CXCL10 blockade, macrophages were incubated with 1.5 ug/ml of anti-CXCL10 (R & D; Cat #: AF466SP) for 45 minutes at 37C.

### *In Vitro* Transwell System for Migration via rmCXCL9 and rmCXCL10

0.15 ug/ml of recombinant mouse protein CXCL9 (Peprotech, Cat #: 250-18) and 0.1 ug/ml of recombinant mouse protein CXCL10 (Peprotech, Cat #: 250-16) pure ligands were plated in the bottom well of a 96 well transwell (Corning, 3387).

## Acknowledgements

The authors thank the patients and families that have donated tissue and blood samples to this study. Additionally, the authors thank the following people for allowing access to their patients for recruitment based on the Institutional Review Board-approved protocol: Drs Alice Fan, Sukhimani Padda, Kavitha Ramchandran, Dimitrios Colevas, Sumit Shah, Maximilian Diehn, Sunil A. Reddy, Sandy Srinivas, and Saad A. Khan.

## Author Contributions

The authors confirm contribution to the paper as follows: study conception and design: YVH, AB, SW, JN, HW, RW, PN, EG, PA, GJB, SMW, HZ, data collection: YVH, YS, HC, BX, ZL, CB, GJB, DL, SW, HZ; analysis and interpretation of results: YVH, HZ, GJB, SMW, draft manuscript preparation: YVH, HZ. All authors reviewed the results and approved the final version of the manuscript.

## Supplemental Material

Supplemental Methods

Supplemental Tables 1A-1D, 2A-2D, 3A-3C Supplemental Figures 1-4

## Non-standard Abbreviations and Acronyms

ICIs: immune checkpoint inhibitors

MRL: Murphy Roths Large

scRNAseq: single-cell RNA-sequencing

scTCRseq: single-cell T-cell receptor sequencing

CITE-seq: cellular indexing of transcriptomes and epitopes

irAEs: immune related adverse events

PD-1: programmed cell death-1

CTLA-4: cytotoxic T-lymphocyte antigen 4

## Supplemental Figure Legends

**Supplemental Figure 1.** Immune cell populations in the hearts of *MRL* and *MRL/Pdcd1-/-* mice. **A)** Quantification of CD45+ level gene expression in all cardiac immune cells. **B)** Feature plots of canonical genes in macrophages/monocytes in the heart. **C)** Quantification of macrophage/monocyte subsets. **D)** UMAP of the CD3+ T-cell level. **E)** Feature plots of T-cell markers. **F)** Quantification of T-cell subsets in the heart. **G)** Feature plots of CD8+ and CD4+ markers. **H)** Quantification of CD8+ T-cell subsets in the heart.

**Supplemental Figure 2.** CellphoneDB analysis of cell-cell interactions in *MRL* and *MRL/Pdcd1-/-* mice. **A)** CXCL signaling pathway network between CD8+ T-cell subsets in the blood and monocyte/macrophage subsets in the heart. **B)** Comparison of CXCL signaling between CD8+ T-cell subsets in the blood and monocyte/macrophage subsets in the heart in *MRL* and *MRL/Pdcd1-/-* mice. **C)** CXCL9-CXCR3 and CXCL10-CXCR3 networks between CD8+ T-cell subsets in the blood and monocyte/macrophage subsets in the heart in *MRL* and *MRL/Pdcd1-/-* mice.

**Supplemental Figure 3.** Immune cell populations in the hearts of *MRL/Pdcd1-/-* mice treated with liposomal control or liposomal clodronate. **A)** Feature plots of canonical cell markers at the CD45+ level. **B)** Quantification of CD45+ immune cell subpopulations. **C)** Feature plots of macrophage cell markers. **D)** Flow gating for circulating macrophage/monocytes for CCR2+ marker. **E)** Quantification of macrophage subpopulations in the heart. **F)** UMAP of T-cell subpopulations. **G)** Feature plots of T-cell subpopulation markers. **H)** Quantification of CD4+ and CD8+ T-cells in the heart. **I)** Feature plots of CD8+ T-cell subpopulation markers. **J)** Quantification of CD8+ T-cell subpopulations.

**Supplemental Figure 4.** Establishment of a novel pharmacological model of ICI myocarditis and immune cell population analysis of the hearts of MRL mice treated with anti-PD1/anti-CTLA-4 immunotherapy and IgG control or anti-CXCR3. **A)** Flow chart of immunotherapy treatment for establishment of pharmacological model of ICI myocarditis. **B)** Flow gating for CD8+ T-cells and macrophages in the hearts of *MRL* mice treated with IgG control or immunotherapy. **C)** Quantification of CD8+ T-cells and macrophages in the hearts of the pharmacological mouse model of ICI myocarditis. **D)** Histology of the hearts of *MRL* mice treated with IgG control or anti-PD1/anti-CTLA-4 immunotherapy. **E)** Quantification of immune cell subsets at the CD45+ level in mice treated with IgG only, immunotherapy only, or immunotherapy and anti-CXCR3. **F)** Quantification of immune cell subsets at the CD3+ level in mice treated with IgG only, immunotherapy only, or immunotherapy and anti-CXCR3. **G)** Quantification of immune cell subsets at the CD8+ level in mice treated with IgG only, immunotherapy only, or immunotherapy and anti-CXCR3.

