## Supplemental Methods for "A Novel Therapeutic Approach using CXCR3 Blockade to Treat Immune Checkpoint Inhibitor-mediated Myocarditis"

Mice

*Genetic PD-1 Deletion Model*

For our time point experiments, we used Murphy Roths Large (MRL) mice as our control population and a genetic mouse model of PD-1 deletion on the MRL background (*MRL/Pdcd1-/-*)^13,47^ for our experimental myocarditis model. *MRL/Pdcd1-/-* mice were obtained from RIKEN (Ibaraki, Japan). MRL mice (strain #000485) were obtained from The Jackson Laboratory (Bar Harbor, Maine). These mice develop spontaneous myocarditis around 4 weeks of age with 70% penetrance.

*Pharmacological Model*

MRL mice were injected intraperitoneally with 400 ug of anti-PD1 monoclonal antibody and 400 ug of anti-CTLA4 monoclonal antibody (BioXCell) or 800 ug of IgG control antibody twice a week starting at 6-7 weeks of age. After six doses, mice were sacrificed.

All animal studies were conducted in accordance with institutional guidelines. All mice were housed in the SIM1 Clean Barrier Facility in the Lokey Stem Cell Research Building at Stanford University, and experiments were conducted in accordance with APLAC protocols.

Sequential Single-Cell Analysis of MRL and *MRL/Pdcd1-/-* PBMCs and Heart Immune Cells at 1 week, 2 weeks, and 4 weeks

MRL and *MRL/Pdcd1-/-* mice were sacrificed at the ages of 1 week, 2 weeks, and 4 weeks via decapitation for 1 week mice and CO2 inhalation and cervical dislocation for the other time points. Blood was collected via cardiac puncture, and both blood and heart were processed to isolate immune cells. See *Heart, Blood, and Peritoneal Macrophage Processing* and *Single-Cell Sequencing* section for further downstream details.

Clodronate-mediated Macrophage Depletion in *MRL/Pdcd1-/-* Mice

*MRL/Pdcd1-/-* mice were injected intraperitoneally with 100 ul of 5 mg/ml control PBS liposomes or liposomal clodronate^48,49^ (LIPOSOMA) every 2 days starting at 2 weeks of age to deplete macrophages. Mice were sacrificed at 4 weeks of age and harvested for heart and blood immune cells. Heart samples were processed for single-cell library creation. Blood samples were analyzed via flow cytometry. See *Heart, Blood, and Peritoneal Macrophage Processing, Histology, Flow Cytometry,* and *Single-Cell Sequencing* section for further downstream details.

*In Vivo* CXCR3 Blockade in MRL Mice treated with anti-PD1 and anti-CTLA4

*Immunotherapy*

MRL mice were injected intraperitoneally with 400 ug of anti-PD1 (BioXCell; Cat#: BE0146) monoclonal antibody and 400 ug of anti-CTLA4 monoclonal antibody (BioXCell; Cat#: BE0164) or associated IgG control antibody twice a week starting at 6-7 weeks of age.

*CXCR3 mAb*

Anti-CXCR3 monoclonal antibody (BioXCell; Cat#: BE0249) or IgG control (BioXCell; Cat#: BE0091) was administered at a dosage of 200 ug, starting two days before immunotherapy treatment and then twice a week at the same time as anti-PD1/anti-CTLA4 treatment. After six doses of immunotherapy, mice were sacrificed.

See *Heart, Blood, and Peritoneal Macrophage Processing, Histology,* and *Single-Cell Sequencing* section for further downstream details

*In Vitro* Transwell System for CXCR3, CXCL9, CXCL10 Blockade

A 24 well plate was coated with 1 ug/ml of anti-CD3 and 3 ug/ml of anti-CD28 (Thermofisher) and incubated at 37C, 5% CO2 for 1-2 hours. *MRL/Pdcd1-/-* heart single cell suspensions were then plated in RPMI + 10% FBS + 10 mM HEPES + 70uM beta-mercaptoethanol + 1% Pen/Strep media supplemented with 20 ng/ml of IL-2. Cells were split every 2-3 days and by 6-7 days for differentiation into CD8+ T-cells^50^.

After a week, heart immune cells were detached via pipetting and isolated for CD45+ cells using the Miltenyi MACS separator CD45 microbeads for mouse (Cat# 130-052-301). For the CXCR3 blockade, CD45+ cells were incubated with 10 ug/ml of anti-CXCR3 (BioXCell) or IgG control (BioXCell) for 30 minutes at 37C.

Peritoneal macrophages from *MRL/Pdcd1-/-* mice were plated in the bottom well of a 96 well transwell (Corning, 3387). For the CXCL9 blockade, macrophages were incubated with 3 ug/ml of anti-CXCL9 (R&D; Cat #: AF492SP) for 45 minutes at 37C. For the CXCL10 blockade, macrophages were incubated with 1.5 ug/ml of anti-CXCL10 (R&D; Cat #: AF466SP) for 45 minutes at 37C. After incubation with antibodies, the isolated CD45 cells from heart PBMC were added to the top insert of the transwell and the assay was left in the incubator for 1.5 hours for migration to occur. All cells from the bottom well were detached via 6 mM of EDTA added to the bottom well and left in the incubator for 10 minutes.

See *Flow Cytometry* for further downstream details.

*In Vitro* Transwell System for Migration via rmCXCL9 and rmCXCL10

A 24 well plate was coated with 1 ug/ml of anti-CD3 and 3 ug/ml of anti-CD28 (Thermofisher) and incubated at 37C, 5% CO2 for 1-2 hours. *MRL/Pdcd1-/-* heart PBMCs were then plated in RPMI + 10% FBS + 10 mM HEPES + 70uM beta-mercaptoethanol + 1% Pen/Strep media supplemented with 20 ng/ml of IL-2. Cells were split every 2-3 days and by 6-7 days for differentiation into CD8+ T-cells. After a week, heart immune cells were detached via pipetting and isolated for CD45+ cells using the Miltenyi MACS separator CD45 microbeads for mouse (Cat# 130-052-301). 0.15 ug/ml of recombinant mouse protein CXCL9 (Peprotech, Cat #: 250-18) and 0.1 ug/ml of recombinant mouse protein CXCL10 (Peprotech, Cat #: 250-16) pure ligands were plated in the bottom well of a 96 well transwell (Corning, 3387). Isolated CD45 cells from heart PBMC were added to the top insert of the transwell and the assay was left in the incubator for 1.5 hours for migration to occur. All cells from the bottom well were detached via 6 mM of EDTA added to the bottom well and left in the incubator for 10 minutes.

See *Flow Cytometry* for further downstream details.

Histology

After sacrifice, the middle of the heart was cut and sent to HistoTec for paraffin block creation, histology processing and H&E staining. H&E slides were viewed under the Keyence microscope. Quantification was conducted on 10x images. Three pictures per heart were taken at the 10x magnification and quantified in an automated fashion via a script written for ImageJ. The total area of purple nuclei in the image was calculated to quantify the purple-stained immune cells as well as nuclei in cardiomyocytes, and this number was then divided by the total area of the heart. A correction factor was subtracted from this percentage to account for the cardiomyocyte nuclei included in the myocarditis area quantification. The percentage of myocarditis was then averaged among the three photos taken for each sample.

Flow Cytometry

*Macrophage Depletion via Liposomal Clodronate*

Blood PBMCs were processed for flow cytometry by staining with the Live/Dead Fixable Zombie Aqua dye from Biolegend for 30 minutes, washing, then blocking with the Biolegend TruStain PLUS (mouse) Fc block for 10 minutes, and then incubating with the following antibodies: CD45 (Invitrogen, 11-0451-82), Ly6g (Biolegend 127627), F4/80 (Biolegend 123111), CD8 (Biolegend, 100729), CD14 (Biolegend 123309), and CCR2 (Biolegend 150621).

*In Vivo CXCR3 Blockade and in Vitro Transwell Assay*

Cells were processed for flow cytometry by staining with the Live/Dead Fixable Zombie Aqua dye from Biolegend for 30 minutes, washing, then blocking with the Biolegend TruStain PLUS (mouse) Fc block for 10 minutes, and then incubating with the following antibodies: CD45 (Invitrogen, 11-0451-82), CD3 (Biolegend 100235), F4/80 (Biolegend 123147), CD8 (Biolegend, 100729), CXCR3 (Biolegend 126533). All cell populations were analyzed via flow cytometry by the Quanteon machine in the Stanford Beckman FACS facility.

Heart, Blood, and Peritoneal Macrophage Processing

Blood was harvested from mice via cardiac puncture after cervical dislocation and incubated in ACK Lysing Buffer (Quality Biological) for 5 minutes. After 5 minutes, 5 ml of RPMI + 10% FBS + 1% Penicillin-Streptomycin was added to quench the lysing. It was then filtered through a 40 um filter. The entire volume was centrifuged at 400g for 5 minutes at 4C. The supernatant was aspirated, and the cells were resuspended in 1 ml of RPMI + 40% FBS + 10% DMSO and stored in a cryogenic tube for liquid nitrogen storage. 10 ul were used for cell counting by mixing with an equal volume of trypan blue and using the Countess 3 machine.

The middle of the heart was cut and sent for histology. The top and bottom sections of the heart tissue were mechanically dissociated using surgical scissors in a buffer of 1 ml of collagenase mixture (10 mg/ml of collagenase A + 10 mg/ml of collagenase B (Sigma) in HBSS+/+ with 40% FBS) and incubated at 37C with shaking for 10 minutes. After digestion, heart sample mixtures were filtered through a 100 um filter. Collected cells were washed with 7 ml of RPMI + 10% FBS + 1% Penicillin-Streptomycin and centrifuged at 400g for 5 minutes at 4C. The supernatant was aspirated, and the cells were resuspended in 1 ml of RPMI + 40% FBS + 10% DMSO and stored in a cryogenic tube for long-term liquid nitrogen storage. 10 ul were used for cell counting by mixing with an equal volume of trypan blue and using the Countess 3 machine.

For peritoneal macrophage isolation^51^, mice were injected intraperitoneally with 500 ul of thioglycolate broth. After 4 days, mice were sacrificed via CO2 inhalation and cervical dislocation. Using forceps and scissors, the skin of the abdomen was peeled back to expose the intact wall of the peritoneum. 5 mL of RPMI + 10% FBS + 1% Penicillin-Streptomycin was injected into the peritoneum using a 20G needle. The peritoneum was shaken gently, and afterwards, fluid was aspirated from the peritoneum using the same needle and syringe. Cell suspensions were centrifuged at 400 g for 5 minutes at 4C to pellet cells. The media was aspirated, and the pellet was resuspended in 10 ml of RPMI + 10% FBS + 1% Penicillin-Streptomycin and filtered through a 40 um filter. Cells were again centrifuged at 400 g for 5 minutes at 4C. The supernatant was aspirated, and the remaining cell pellet was resuspended in 1 ml of RPMI + 40% FBS + 10% DMSO and stored in a cryogenic tube. 10 ul were used for cell counting by mixing with an equal volume of trypan blue and using the Countess 3 machine.

All cell suspensions in RPMI + 40% FBS + 10% DMSO were stored at -80C for 24 hours before being moved the liquid nitrogen.

Single-Cell Sequencing

The base and apex of the heart were processed for PBMCs and isolated for CD45+ immune cells that were sent to Medgenome for single-cell sequencing. CD45+ immune cells were isolated from heart PBMCs via Miltenyi MACS magnet and CD45+ Microbeads.

Samples were incubated with 50cc anti-mouse TruStain FcX PLUS (CD16/32) antibody (BioLegend) for 10 minutes on ice and then with 50cc of an antibody master mix containing TotalSeq anti-mouse Hashtag antibody, TotalSeq anti-mouse/human CD44 antibody (BioLegend, 103063, dilution 1:50) and TotalSeq anti-mouse/human CD62L antibody (BioLegend, 104455, dilution 1:50) for 20 minutes on ice.

Frozen PBMCs were removed from liquid nitrogen and resuspended in warm RPMI media + 10% FBS (@ 37ºC) by adding 1 mL, 3 mL, 4 mL, 8 mL 9 mL media at 30 second intervals. Blood cells were washed and passed through a 40 um filter, centrifuged @ 400g for 5 min resuspended in PBS with 50 ul 1% BSA and Biolegend Fc Receptor Blocking Solution (Mouse TruStain PLUS Fc block Cat. No. 156603) for 10 minutes. Heart cells were passed through a 40 um filter and then through a MACS LS column with CD45 microbeads (Cat. No. 130-052-301, Milltenyi, Bergisch Gladbach, North Rhine-Westphalia, Germany) to enrich for immune cells prior to undergoing the above blocking process. 50 uL of antibody mastermix containing Biolegend TotalSeq-C antibodies for staining (TotalSeq™-C0301 anti-mouse Hashtag 1 Antibody Cat. #. 155861; TotalSeq™-C0302 antimouse Hashtag 2 Antibody Cat. #. 155863; TotalSeq™-C0303 anti-mouse Hashtag 3 Antibody Cat. #. 155865; TotalSeq™-C0304 anti-mouse Hashtag 4 Antibody Cat. #. 155867; TotalSeq™-C0305 anti-mouse Hashtag 5 Antibody Cat. #. 155869; TotalSeq™-C0306 anti-mouse Hashtag 6 Antibody Cat. #. 155871; TotalSeq™-C0307 anti-mouse Hashtag 7 Antibody Cat. #. 155873; TotalSeq™- C0308 anti-mouse Hashtag 8 Antibody Cat. #. 155875; TotalSeq™-C0073 anti-mouse/human CD44 Antibody Cat. #. 103063; TotalSeq™-C0112 anti-mouse CD62L Antibody Cat. #. 104455) at the manufacture-recommended concentrations were then added to each sample. After incubation, cells were washed 2x by resuspending in 1.5 mL PBS + 1% RNase-free BSA (MACS BSA Stock Solution No. 130-091-376) and then 1 mL PBS + 0.04% BSA. 10 uL of cell suspension was then taken for counting and appropriate volume of PSA + 0.04% BSA was added (200-500 uL) to aim for final target concentration of 1-1.5 million cells/mL). Cells were then loaded into Chromium Next GEM Chip K (10x PN 2000182).

Samples were targeted for 10,000 cells and libraries generated via the single-cell 5’ RNA and TCR 10x Chromium system. Libraries were sequenced by Medgenome with 20,000 pair reads for the gene expression library and 5,000 pair reads for the TCR and surface protein libraries.

*Library Preparation for scRNAseq*

Library preparation was completed according to 10x’s manufacturing protocols for the Chromium Next GEM Single Cell V(D)J Reagent Kits v1.1 and Chromium Next GEM Single Cell V(D)J Reagent Kits v2. Isolated cDNA was amplified, and then cDNA was allocated for TCR enrichment, feature barcode library generation, and 5’ cDNA library generation. cDNA and library quality were evaluated using the high sensitivity DNA kit on the Agilent 2100 Bioanalyzer.

*Sequencing of scRNAseq libraries*

Sequencing of all 10x Genomics libraries was performed through Medgenome (Foster City, California). 5’ library, TCR library, and feature barcoding libraries were all sequenced on the Illumina NovaSeq 6000 instrument using 200 cycle kits and run with 2x100bp for a goal of 20,000 paired reads/cell, 5,000 paired reads/cell, and 5,000 paired reads/cell.

*Single Cell RNA-seq Data Pre-Processing*

Raw single-cell RNA-seq data was processed using 10x Genomics Cell Ranger (6.0.2) to demultiplex the FASTQ reads in order to align them to the mouse reference genome (refdata-gex-mm10-2020-A, provided through the 10x Genomics website) and to count the unique molecular identifier (UMI) (through the “cellranger count” function). Individual sample gene expression matrices were loaded into Seurat v4.0.5 R package (https://satijalab.org/seurat/) for further analysis.

To filter out poor quality data from cells likely to be dead/damaged cells, we removed cells with greater than >20% mitochondrial genes present. Additionally, we removed cells for which less than 200 genes and more than 5500 were detected. Doublets from multiplexed samples were removed using duplicate barcoded hashtag counts from cells. After RBC and fibroblast removal, the timepoint sequencing data had ~70,000 immune cells, the macrophage depletion data had ~17,500 immune cells, and the *in vivo* CXCR3 blockade data had ~16,200 immune cells.

*Identification of Cell Clusters*

The count matrix for each sample in the time point experiments was normalized and integrated via the fast mutual nearest neighbors^18^ correction which corrects for batch effects in single-cell data using a version of the mutual nearest neighbors (MNN) method. The count matrix for each sample in the macrophage depletion and *in vivo* CXCR3 blockade experiments was normalized using the ‘SCTransform’ function in Seurat^19^, which reveals sharper biological distinctions with better correction for sequencing depth and batch effects compared to the standard Seurat workflow^19^. All samples were then integrated into a single Seurat object using the reciprocal PCA (RPCA) multi-modal data integration pipeline in Seurat, through the determination of anchors by the mutual neighborhood requirement. To perform clustering analysis, principal components (PCs) used for downstream analyses were selected by ranking the principal components based on the amount of variance added by each additional PC, and an elbow plot was generated to select the number of PCs at the point where additional PCs did not explain additional variance in the data. These PCs were then used for constructing the shared nearest neighbor (SNN) graph or mutual nearest neighbors (MNN) by using ‘FindNeighbors’ function. Louvain clustering algorithm was then used to cluster the cells.

*Annotation of Cell Clusters and Data Visualization*

We utilized canonical markers of different immune cell subtypes and checked whether these well-studied genes were the top differentially expressed genes of each queried cluster. The uniform manifold approximation and projection (UMAP) was applied to visualize the single cell transcriptional profile in a two-dimensional space based on the SNN graph. Feature plots and violin plots were generated using Seurat’s standardized code based on ggplot2 (v3.2.1, R package). Subsets of CD45+ cells (CD3+, CD8+, and macrophage/monocyte subsets) were analyzed by subsetting CD45+ cell clusters on basis of canonical markers and re-applying the clustering algorithm.

*Differential Expression Analysis*

Differential expression tests were performed in the Seurat R package^45,46^ using FindMarkers() and FindAllMarkers() methods, based on the non-parametric Wilcoxon rank sum test with a logfc.threshold set to the default of 0.25 (limiting testing to genes which show, on average, at least 0.25 difference on the log-scale between two groups of cells). Multiple testing correction was performed based on the Bonferroni correction using all features in the dataset to give an adjusted p-value. Resulting genes with adjusted p-values above the significant threshold of > 0.05 were filtered out.

*T Cell Receptor Data Analysis*

Raw T cell receptor (TCR) sequencing data was processed with 10x Genomics Cell Ranger (v6.0.2). The algorithm aligned FASTQ reads to human GRCh38 V(D)J reference genome (v4.0.0, from 10X Genomics) by using the “cellranger vdj” function, resulting in the assembly of V(D)J sequences and clonotypes. The filtered contig annotations, which contained high-level annotations of each high-confident cellular contig, was used. Using Python, the clonotype ID and cell barcode IDs within each unique sample were paired and inputted into Seurat^45,46^ for visualization of individual T-cell clonotypes on the UMAPs. An identical pipeline was applied for the mouse single-cell TCR-seq data.

*Ligand-Receptor Expression and Cell Interactions*

We examined the interaction of CD8+ T-cells with our macrophages/monocytes in our time point mouse scRNAseq data based on the expression level of ligand-receptor pairs using the CellChatDB^22^ database.
