## Supplementary figures and images for "A Novel Therapeutic Approach using CXCR3 Blockade to Treat Immune Checkpoint Inhibitor-mediated Myocarditis"

### Supplemental Figure 1

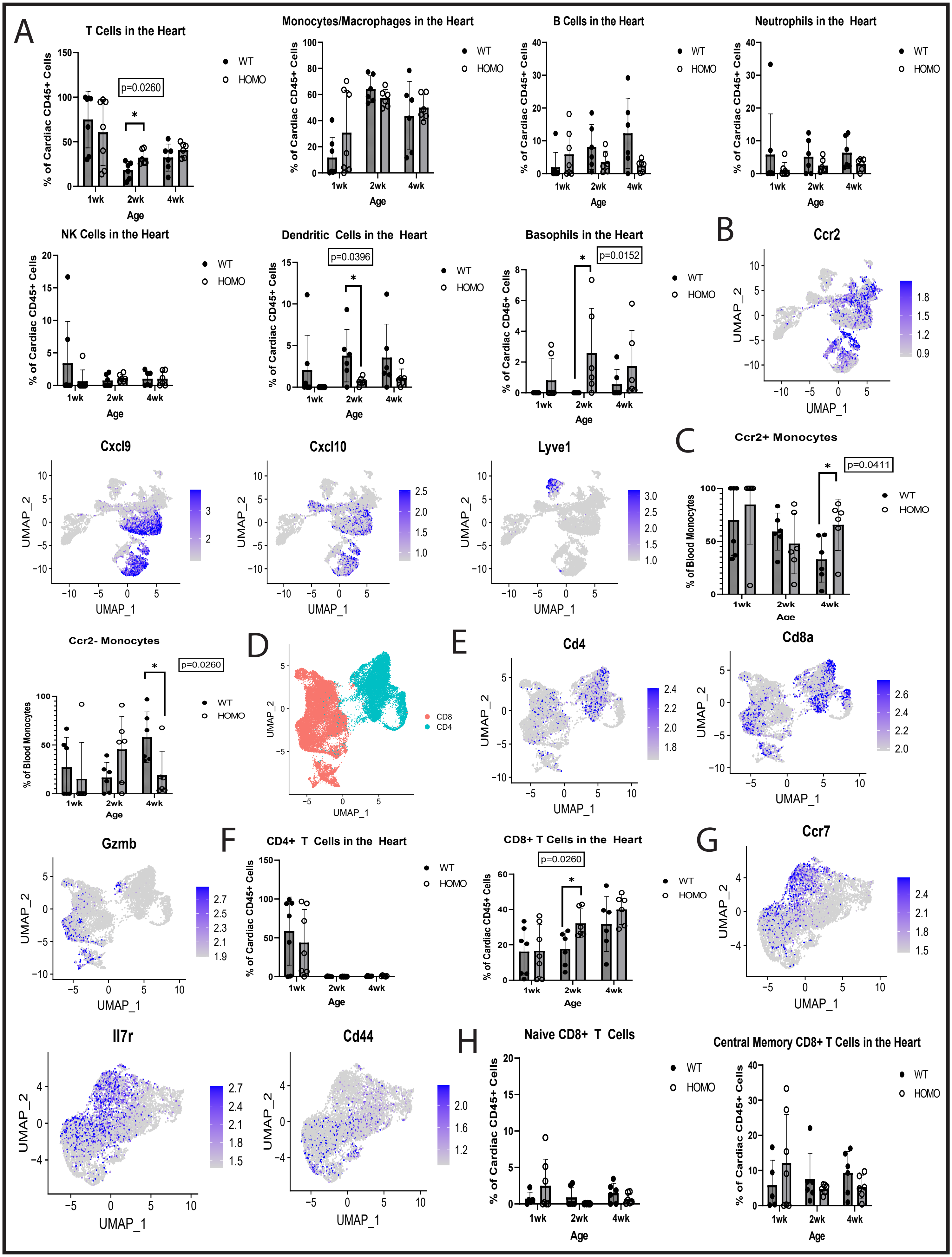

### Supplemental Figure 2

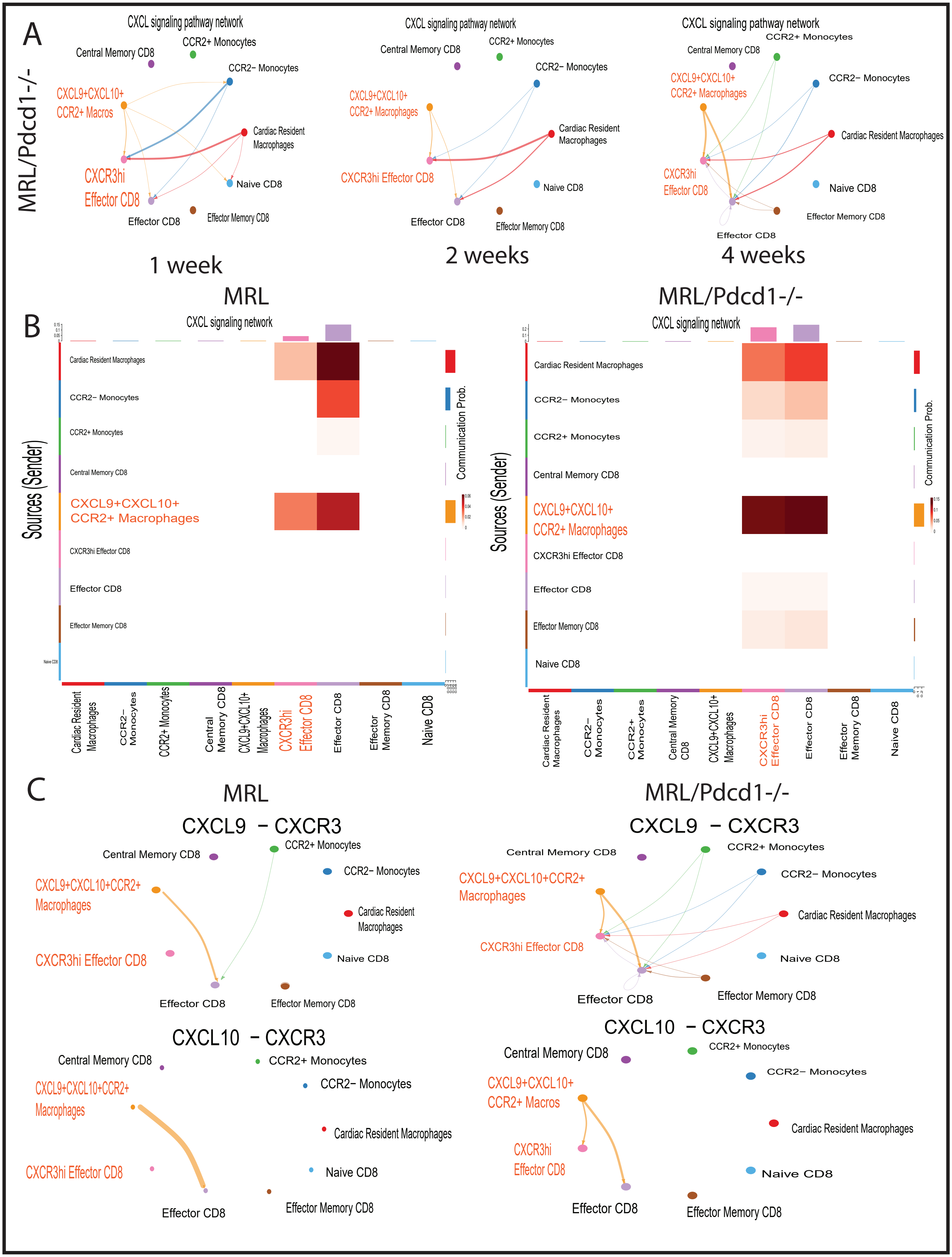

### Supplemental Figure 3

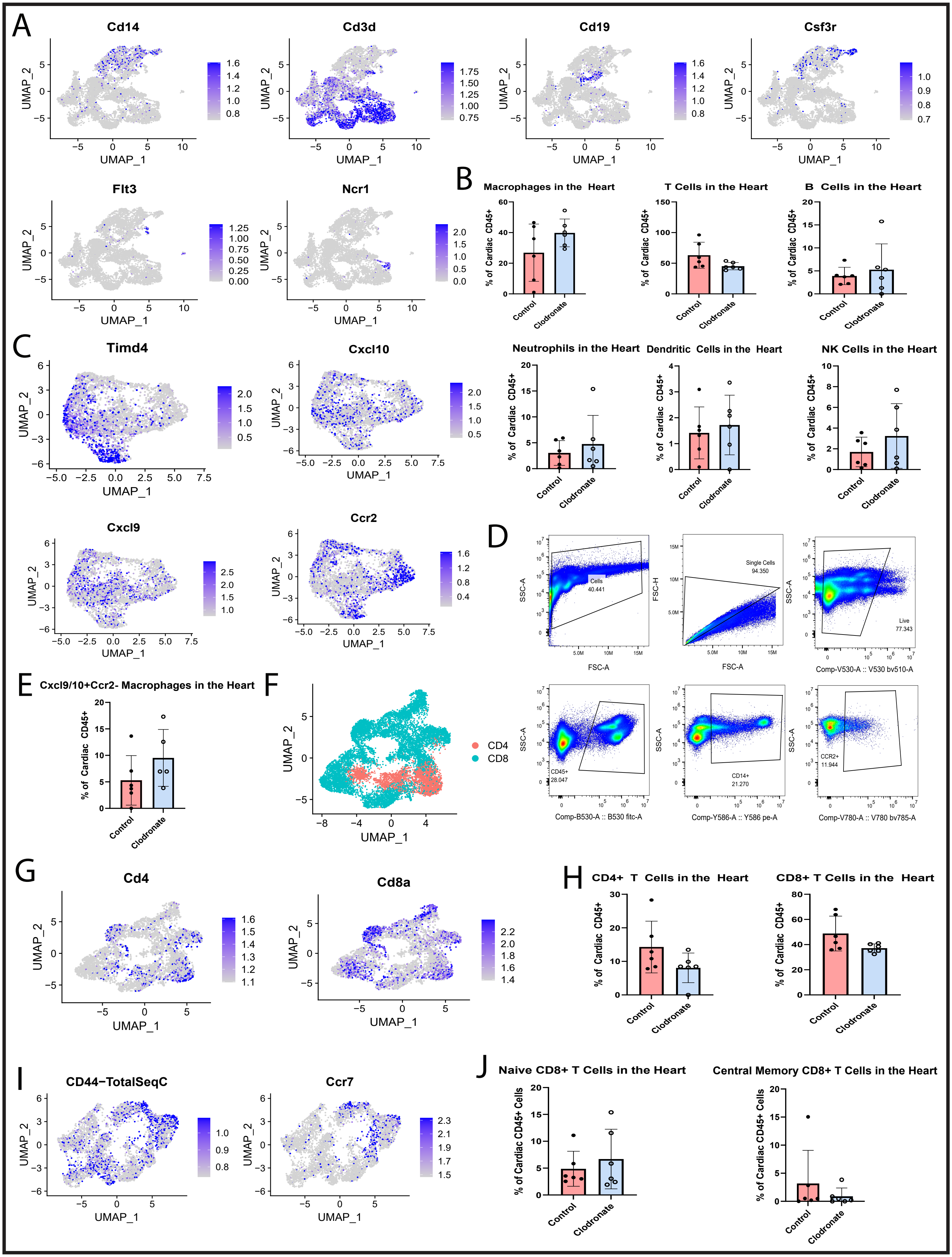

### Supplemental Figure 4

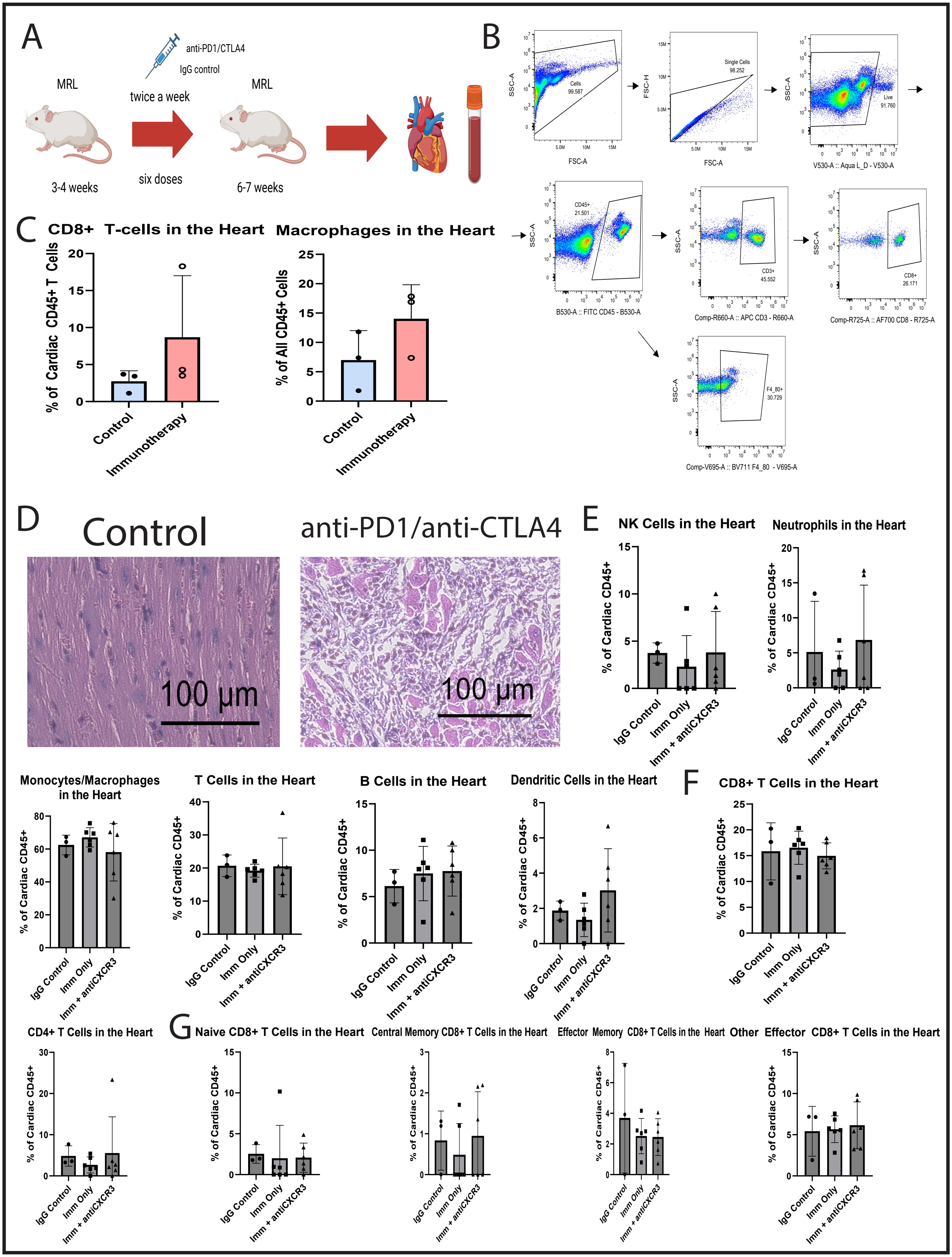
